# Disentangling a Complex Response in Cell Reprogramming and Probing the Waddington Landscape by Automatic Construction of Petri Nets

**DOI:** 10.1101/599191

**Authors:** Viktoria Rätzel, Britta Werthmann, Markus Haas, Jan Strube, Wolfgang Marwan

## Abstract

We analyzed the developmental switch to sporulation of a multinucleate *Physarum polycephalum* plasmodial cell, a complex response to phytochrome photoreceptor activation. Automatic construction of Petri nets from trajectories of differential gene expression in single cells revealed alternative, genotype-dependent interconnected developmental routes and identified metastable states, commitment points, and subsequent irreversible steps together with molecular signatures associated with cell fate decision and differentiation. Formation of transition-invariants in mutants that are locked in a proliferative state is remarkable considering the view that oncogenic alterations may cause the formation of cancer attractors. We conclude that the Petri net approach is useful to probe the Waddington landscape of cellular reprogramming, to disentangle developmental routes for the reconstruction of the gene regulatory network, and to understand how genetic alterations or physiological conditions reshape the landscape eventually creating new basins of attraction. Unraveling the complexity of pathogenesis, disease progression, and drug response or the analysis of attractor landscapes in other complex systems of uncertain structure might be additional fields of application.

## 1 Introduction

Cell differentiation, the formation of functionally specialized cells by selective expression of cell type-specific genes, is deeply rooted in the evolution of eukaryotes (Russo et al. 1999; Schaap et al. 2016). In simple eukaryotes, cells of specialized form and function may develop in temporal order in the course of a life cycle instead of developing concurrently to form a multicellular organism. There is growing evidence that the reprogramming of gene expression during commitment and differentiation occurs with the dynamics of a complex system that switches between stable or meta-stable states of gene expression (Ferrell Jr 2012; Graf and Enver 2009; Huang et al. 2009; Zhou and Huang 2011). Non-linear effects contributing to the dynamic behavior of the system may explain the experimentally observed natural heterogeneity in developmental trajectories of individual mammalian cells (Ferrell Jr 2012; Li et al. 2016) and of cells of simple eukaryotes (Solnica-Krezel et al. 1991; Werthmann and Marwan 2017) within clonal populations. They may also explain the outcome of artificial cell reprogramming experiments as they are performed for the generation of induced pluripotent stem (iPS) cells ((Il Joo et al. 2018) and references therein). As informal intuitive reasoning based on proximate causation is not sufficient to fully understand the behavior of complex regulatory networks ((Tyson et al. 2019; Yildirim and Huang 2018) and references therein), theoretically sound approaches (Chen et al. 2014; Il Joo et al. 2018; Wang et al. 2011; Wu et al. 2017; Zhou et al. 2012) based on a quasi-potential landscape of cellular reprogramming have been developed and were metaphorically illustrated with the help of the epigenetic landscape proposed by Waddington ((Huang et al. 2009; Zhou and Huang 2011) and references therein). In Waddington’s paradigm, the developmental trajectory of a cell proceeds like a marble rolling down a landscape with multiple bifurcating valleys finally ending up in one of separate valleys each of which corresponds to an alternative state of terminal differentiation (Waddington 1957). In a rugged quasi-potential landscape of a dynamic system, a trajectory may take alternative paths (Zhou and Huang 2011), resulting in considerable cell to cell variability.

Individually different developmental pathways experimentally demonstrated in mammalian cells have been explained based on the Waddington landscape (Huang et al. 2007; Zhou et al. 2016) and are evident even at the morphological level in *Physarum polycephalum.* The development of a *P. polycephalum* amoeba into a unicellular, multinucleate plasmodium occurs through intermediate morphological stages with fundamental, discrete differences in the organization of cytoskeleton and nucleus (Solnica-Krezel et al. 1991).

Like amoebae, individual plasmodial cells also take different pathways of differential gene expression to commitment and differentiation (sporulation) (Rätzel and Marwan 2015; Werthmann and Marwan 2017) which involves extensive remodeling of the regulatory network at the transcriptional, translational, and post-translational level (Glöckner and Marwan 2017). Therefore, and because of the possibility to retrieve multiple samples from the same cell (see below), plasmodial cells are ideally suited for the analysis of developmental variability (Rätzel and Marwan 2015).

*P. polycephalum* forms multinucleate giant plasmodial cells that can be grown to arbitrary macroscopic size under controlled axenic conditions. Due to the continuous, vigorous streaming and mixing of the suspending cytoplasm the nuclei of a plasmodial cell display natural synchrony in cell cycle and differentiation (Guttes and Guttes 1961; 1964; Hoffmann et al. 2012; Rusch et al. 1966; Sachsenmaier et al. 1972; Sauer et al. 1969; Starostzik and Marwan 1995a). Developmental synchrony is also evident through the response to threshold stimulation, where the developmental decision to sporulation is all or none for each individual plasmodium (Starostzik and Marwan 1994; 1995a; 1998) and, well after commitment, through the simultaneous conversion of the plasmodial mass into fruiting bodies (Hoffmann et al. 2012; Sauer et al. 1969). Multiple samples simultaneously taken from different sites of the same plasmodium showed no significant differences in the expression patterns of the 35 genes analyzed in this work (Rätzel 2015). Even two plasmodia of different sporulation-deficient genotype, one stimulated and one un-stimulated, self-synchronize their gene expression patterns after plasmodial fusion and the outcome is consistent with the developmental destiny of the cell (Walter et al. 2013).

This natural synchrony and homogeneity of the plasmodium encourages single cell time series analyses by repeatedly taking samples of the same plasmodium after application of a differentiation-inducing stimulus. To set a defined starting point, the differentiation into fruiting bodies (sporulation) can be triggered by a brief pulse of far-red light (Rätzel and Marwan 2015) which is sensed by phytochrome, a light-switchable photoreceptor (Lamparter and Marwan 2001; Schaap et al. 2016; Starostzik and Marwan 1995b). Although previous work has shown that individual plasmodial cells take variable pathways to commitment and differentiation (Rätzel and Marwan 2015), specific main routes which trajectories might follow, have not been identified. These routes however are important to know because averaging over a cell population is presumably inappropriate for a thorough analysis of the molecular architecture of the gene regulatory network.

Following up previous work in which we analyzed the physiological states of a cell on the way to commitment and sporulation (Marwan et al. 2005), we recently found that the states of gene expression discretized by hierarchical clustering can be turned into a Petri net model that predicts single cell trajectories in response to a differentiation-inducing stimulus (Werthmann and Marwan 2017). Because this approach was heuristic and relied on the manual composition of Petri nets, it could be tried on a small data set only.

In order to explore the potential power of Petri nets to model, predict, analyze, and understand the dynamic behavior of individual cells we now developed a method to automatically construct Petri nets from time series data and applied it to a considerably larger data set obtained from wild type and mutants with altered propensity to differentiate.

## 2 Materials and Methods

### 2.1 Strains, growth of cells, sample preparation, and gene expression analyses

Cells of PHO3 (Sujatha et al. 2005), PHO57, PHO64 (Rätzel et al. 2013), and WT31 (originally isolated as CS310) (Starostzik and Marwan 1998) were grown and starved under standard conditions in the dark as described (Rätzel et al. 2013). Starved plasmodial cells (each Petri dish of 9 cm diameter contained one plasmodial cell) were stimulated with a pulse of far-red light (λ≥700 nm, 13 W/m^2^) (Starostzik and Marwan 1998) of the length indicated in Table 1 and returned to the dark at 22°C. Before (dark control; referred to as 0 h sample) and at various time points (2h, 6h, 8h, 11h) after the stimulus pulse, samples from the plasmodium were taken with a spatula and shock frozen in liquid nitrogen for the subsequent isolation of RNA as described (Rätzel and Marwan 2015). Single cell gene expression time series data were obtained by measuring the relative mRNA abundance of a set of 35 genes, differentiation marker and reference genes (Hoffmann et al. 2012) (SI Table 1) by gene expression profiling (GeXP), a multiplex RT-PCR method (Hayashi et al. 2007) as described (Marquardt et al. 2017; Rätzel 2015).

**Table 1.** Comprehensive data set combined from different experiments with mutants and wild type (WT31). Data of wild type cells were taken from (Rätzel and Marwan 2015).

| Data Subset | Strain | Starved for (d) | Far-red (min) | No of Cells | Sporulated | Places | Transitions | Immediate Transitions | T-Invariants |
| --- | --- | --- | --- | --- | --- | --- | --- | --- | --- |
| 1 | PHO3 | 6 | 0 | 5 | y | 13 | 14 | 3 | 0 |
| 2 | PHO3 | 6 | 60 | 8 | n/y | 18 | 25 | 4 | 2 |
| 3 | PHO64 | 6 | 0 | 14 | n | 17 | 28 | 10 | 13 |
| 4 | PHO64 | 6 | 30 | 17 | n | 19 | 46 | 8 | 329 |
| 5 | PHO57 | 6 | 0 | 9 | n | 15 | 18 | 6 | 5 |
| 6 | PHO57 | 6 | 30 or 60 | 13 | n | 22 | 33 | 7 | 13 |
| 7 | WT31 | 7 | 0 | 8 | n | 14 | 11 | 7 | 4 |
| 8 | WT31 | 7 | 2 or 5 | 13 | n | 29 | 39 | 8 | 3 |
| 9 | WT31 | 7 | 2 or 5 | 16 | y | 25 | 36 | 5 | 0 |
| 10 | WT31 | 7 | 20 or 30 | 13 | y | 38 | 42 | 8 | 0 |
| 11 | WT31 | 6 | 5 or 15 | 13 | y | 24 | 35 | 4 | 0 |

### 2.2 Data analysis pipeline and automated generation of Petri nets

To correct for potential variations in the concentration of total RNA and in the efficiency of the RT-PCR reaction, expression data were normalized to the median of all 35 values obtained for each RNA sample. The values were then, for each gene separately, normalized to the geometric mean of all values of the respective gene within the data set.

The resulting normalized data sets were further evaluated with the help of a data analysis pipeline written in R (R Core Team 2016). For a given set of genes, the pipeline determined significantly different clusters by running the *Simprof* algorithm (Clarke et al. 2008) as part of the *clustsig* package (Whitaker and Christman 2014), generated a heatmap using the *heatmap.2* function as part of the *gplots* package (Warnes et al. 2016), performed classical multidimensional scaling analysis based on the euclidean distance (Gower 1966) using the *cmdscale* function of the *stats* package v3.5.1 (R Core Team 2016), and displayed the multidimensional scaling results according to different criteria. Along with other files for analysis and documentation, the pipeline also determined single cell trajectories. Each trajectory consisted of a sequence of states of gene expression in which the plasmodium had been at any sampled time point (0h, 2h, 6h, 8h, 11h) after the start of the experiment. Each state was defined through the identity (ID) number of the *Simprof* cluster to which a particular gene expression pattern had been assigned. These trajectories were then evaluated for the transits between states which a cell underwent in changing from one state to a different next state. For the sake of clarity, we discriminate the terms *transit* and *transition.* The term *transit* refers to the change of a cell between states while the term *transition* refers to the corresponding Petri net node that connects the two places representing two subsequent states, respectively. The R script determined the number of transits that occurred between each pair of different states. This number of transits (events) was stored in the Petri net description vector (PNDV) (SI Fig. 4) for becoming part of the name of the corresponding transition. For details see SI Computational Methods.

The PNDV files exported from R were converted into ANDL files, imported as stochastic Petri nets into Snoopy (Heiner et al. 2012; Rohr et al. 2010), and graphically displayed by employing automatic layout algorithms provided by Snoopy. For further details concerning the user-specified combination of Petri nets and the coloring of transitions according to their occurrence in specific subsets of the data, see SI Computational Methods.

### 2.3 Structural analysis of Petri nets

After manually deleting the stack of unused places (not connected to transitions) which for technical reasons are still present in those Snoopy files that represent only a subset of the data, the Petri net was exported as ANDL file from Snoopy for the subsequent determination of source places, sink places, and T-invariants with the help of the analysis tool Charlie (Heiner et al. 2015). These elements were then colored by loading the result files generated by Charlie into the open Snoopy file that displayed the Petri net graphically, using the *Load node set file* function of Snoopy. A video tutorial on computation and graphical display of T-invariants is given as supplementary file in the electronic version of (Werthmann and Marwan 2017).

## 3 Results

Time series data were obtained by retrieving samples of individual plasmodial cells repeatedly, at different time points after the start of the experiment (Rätzel and Marwan 2015). In cells stimulated with far-red light, one sample (0h sample) was taken immediately before delivery of the stimulus. Cells were returned to the dark and the following samples were taken at 2h, 6h, 8h, and 11h after onset of the far-red pulse in order to cover the period before and after commitment. In time series of dark controls, the light pulse was omitted and the 0h sample refers to the start of the experiment. To analyse the variability and the sporulation-specificity of gene expression patterns data were taken from different experiments with the wild type and from three mutant strains either non-sporulating or with strongly reduced propensity to sporulate (Table 1).

The expression patterns of 35 genes, differentiation marker and reference genes (SI Table 1), were determined by gene expression profiling (GeXP), a multiplex RT-PCR protocol combined with capillary electrophoresis (Hayashi et al. 2007; Hoffmann et al. 2012; Marquardt et al. 2017). Based on their similarity to genes in the Uniprot database, the marker genes were chosen for being involved in various cellular functions (signal transduction, cell cycle regulation, transcriptional control, RNA-binding *etc.* (Hoffmann et al. 2012)).

To correct for potential differences in the total RNA concentration and in the efficiency of the RT-PCR reaction between samples, the expression values of each sample were normalized to the median of all 35 values obtained for each respective RNA sample. We observed that although the *ribB* mRNA (ribB) was not differentially regulated by far-red light and remained constant during the measured time intervals, the variation of ribB abundance was considerably higher between individual plasmodia than between the samples taken from the same plasmodium (not shown). Hence, we determined the data quality by dividing the 5 values for ribB of each time series and plotted the frequency distribution of the log2 of all of these ratios (SI Fig. 1). From 645 values analyzed, 96.3% were between −1 and 1 with the by far most abundant class at zero, validating the method of data normalization and indicating that the errors in measurement were small as compared to the observed differences in gene expression patterns.

In order to make the degree of light regulation comparable between genes while maintaining individual differences in gene-specific expression patterns between individual plasmodial cells and between wild type and mutants, the expression values for each gene were subsequently normalized to the geometric mean of all expression values of each respective gene within the considered data set. From the resulting heat map of the complete data set (SI Fig. 2), 19 strongly light-regulated genes *psgA, pldC, pldB, pikB, nhpA, pumA, meiB, cudA* (down-regulated) and *hstA, arpA, pptA, ehdA, cdcA, pakA, gapA, pldA, ligA, pwiA*, and *rgsA* (up-regulated) were selected for further analysis. The expression patterns of the 19 strongly differentially regulated genes are displayed in the heat maps of Fig. 1 to reveal differences between the non-sporulating mutants and the wild type.

**Fig. 1.**
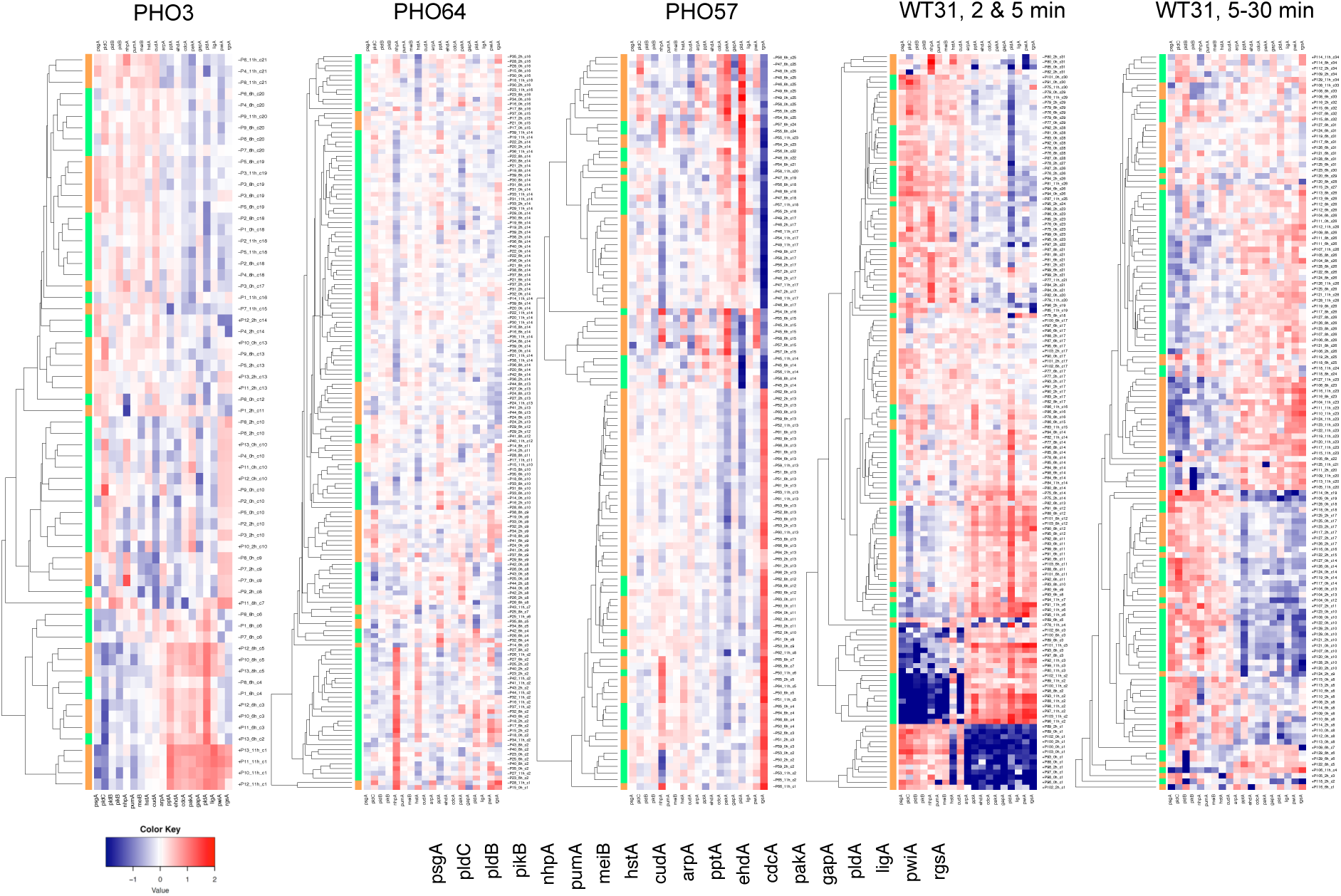
Expression patterns of differentially regulated genes in mutants with reduced (PHO3) or strongly reduced (PHO64, PHO57) propensity to sporulate compared to the wild type (WT31). Expression data of cells exposed to a far-red light pulse and of not exposed cells (dark controls; see Table 1 for details) were pooled, clustered for each indicated group of cells separately, and displayed as heat maps. Side bars indicate Simprof cluster 10 of x-fold gene-specific differences in relative mRNA abundance. The order of the genes for which the data are displayed in each of the heat maps is shown at the bottom.

For further analysis, all expression data (Table 1) were combined into one comprehensive data set. Significantly different clusters of expression were identified with the help of the *Simprof* algorithm (Clarke et al. 2008; Whitaker and Christman 2014). The cluster identity number as determined by the Simprof analysis algorithm was assigned to each expression pattern *i.e.* to each set of expression values of the 19 genes for a given plasmodium at the respective time point (see SI Fig. 3 for a heat map where the significant clusters are indicated by the side bar). A heat map showing the data set in the form the gene expression time series for each individual plasmodium is part of SI File Package 1 in the Supplemental Information available in the electronic version of this article.

**Table 2.** Single cell trajectories of cells stimulated at threshold. Class numbers refer to the data subsets listed in Table 1.

| Strain | Starv | FR | Spo | Class | P | Trajectory |  |  |  |  | P |
| --- | --- | --- | --- | --- | --- | --- | --- | --- | --- | --- | --- |
|  |  |  |  |  |  | 0h | 2h | 6h | 8h | 11h |  |
| WT31 | 7 | 0 | 0 | 7 | P67 | 105 | 103 | 103 | 103 | 103 | P67 |
| WT31 | 7 | 0 | 0 | 7 | P68 | 91 | 91 | 91 | 91 | 91 | P68 |
| WT31 | 7 | 0 | 0 | 7 | P69 | 89 | 98 | 89 | 89 | 97 | P69 |
| WT31 | 7 | 0 | 0 | 7 | P70 | 100 | 100 | 99 | 100 | 100 | P70 |
| WT31 | 7 | 0 | 0 | 7 | P71 | 97 | 104 | 104 | 104 | 89 | P71 |
| WT31 | 7 | 0 | 0 | 7 | P72 | 105 | 105 | 105 | 94 | 105 | P72 |
| WT31 | 7 | 0 | 0 | 7 | P73 | 96 | 96 | 96 | 96 | 96 | P73 |
| WT31 | 7 | 0 | 0 | 7 | P74 | 101 | 101 | 101 | 101 | 92 | P74 |
| WT31 | 7 | 2 | 0 | 8 | P75 | 91 | 26 | 26 | 55 | 95 | P75 |
| WT31 | 7 | 2 | 0 | 8 | P76 | 91 | 91 | 98 | 101 | 98 | P76 |
| WT31 | 7 | 2 | 0 | 8 | P77 | 105 | 89 | 89 | 112 | 88 | P77 |
| WT31 | 7 | 2 | 0 | 8 | P78 | 101 | 93 | 112 | 111 | 4 | P78 |
| WT31 | 7 | 2 | 0 | 8 | P79 | 98 | 98 | 105 | 98 | 90 | P79 |
| WT31 | 7 | 5 | 0 | 8 | P80 | 11 | 11 | 36 | 29 | 8 | P80 |
| WT31 | 7 | 5 | 0 | 8 | P81 | 104 | 106 | 106 | 106 | 91 | P81 |
| WT31 | 7 | 5 | 0 | 8 | P82 | 106 | 13 | 86 | 89 | 112 | P82 |
| WT31 | 7 | 5 | 0 | 8 | P83 | 102 | 89 | 44 | 38 | 110 | P83 |
| WT31 | 7 | 5 | 0 | 8 | P84 | 106 | 106 | 29 | 29 | 29 | P84 |
| WT31 | 7 | 5 | 0 | 8 | P85 | 11 | 91 | 29 | 112 | 114 | P85 |
| WT31 | 7 | 5 | 0 | 8 | P86 | 91 | 91 | 110 | 111 | 111 | P86 |
| WT31 | 7 | 5 | 0 | 8 | P87 | 101 | 91 | 89 | 97 | 90 | P87 |
| WT31 | 7 | 2 | 1 | 9 | P88 | 2 | 87 | 43 | 47 | 1 | P88 |
| WT31 | 7 | 2 | 1 | 9 | P89 | 2 | 2 | 54 | 7 | 1 | P89 |
| WT31 | 7 | 2 | 1 | 9 | P90 | 89 | 89 | 43 | 47 | 22 | P90 |
| WT31 | 7 | 2 | 1 | 9 | P91 | 92 | 89 | 43 | 47 | 22 | P91 |
| WT31 | 7 | 2 | 1 | 9 | P92 | 101 | 104 | 44 | 44 | 7 | P92 |
| WT31 | 7 | 2 | 1 | 9 | P93 | 2 | 89 | 37 | 7 | 1 | P93 |
| WT31 | 7 | 5 | 1 | 9 | P94 | 91 | 91 | 91 | 112 | 40 | P94 |
| WT31 | 7 | 5 | 1 | 9 | P95 | 91 | 91 | 32 | 46 | 22 | P95 |
| WT31 | 7 | 5 | 1 | 9 | P96 | 2 | 114 | 87 | 33 | 1 | P96 |
| WT31 | 7 | 5 | 1 | 9 | P97 | 2 | 64 | 87 | 7 | 1 | P97 |
| WT31 | 7 | 5 | 1 | 9 | P98 | 2 | 2 | 29 | 1 | 1 | P98 |
| WT31 | 7 | 5 | 1 | 9 | P99 | 91 | 28 | 28 | 44 | 7 | P99 |
| WT31 | 7 | 5 | 1 | 9 | P100 | 2 | 2 | 87 | 7 | 1 | P100 |
| WT31 | 7 | 5 | 1 | 9 | P101 | 92 | 89 | 44 | 47 | 7 | P101 |
| WT31 | 7 | 5 | 1 | 9 | P102 | 2 | 2 | 89 | 7 | 1 | P102 |
| WT31 | 7 | 5 | 1 | 9 | P103 | 2 | 89 | 43 | 47 | 1 | P103 |

To further visualize differences in expression patterns between mutants and wild-type and between successive time points for individual cells, we performed multidimensional scaling (MDS) analysis of the gene expression data in two dimensions (k=2; and displayed the results using R (Gower 1966; R Core Team 2016). There were clear differences between wild type and non-sporulating mutants in the localization of the data points within the MDS plot (Fig. 2). While there were some differences in the relative localization of the data points of unstimulated as compared to far-red light-stimulated PHO3 cells, some of which sporulated in response to the stimulus (Fig. 2A,B), there were minor, if any differences between unstimulated and light-stimulated cells in the strictly non-sporulating mutants PHO64 and PHO57 (Fig. 2C-F). Clearly, PHO57 cells formed two sub-populations in both, in the light stimulated sample and in the dark controls (Fig. 2E,F). In contrast to the mutants, there was a clear response to far-red light in sporulating and even in not sporulating wild type cells (see below) as compared to the dark controls (Fig. 2G-K). To discriminate between expression patterns caused by light stimulation *per se* and those associated with light-induced sporulation, we exposed wild type plasmodia to a pulse of far-red light which was so short, that only some of the plasmodial cells sporulated. Even in response to weak stimulation, sporulation of individual plasmodial cells was still all or none, *i.e.* in sporulating cells the entire cytoplasmic mass differentiated into fruiting bodies while stimulated but not sporulating cells behaved equally homogenously in that not even a single fruiting body was formed (Rätzel and Marwan 2015; Starostzik and Marwan 1995b). Those wild type plasmodial cells that were not sporulating in response to the weak stimulus still displayed a changed gene expression pattern as compared to the dark controls (Fig. 2G,H) while the response of sporulating cells within the same experiment was much more pronounced with data points moving with time from upper right to lower left of the plot (Fig. 21). Nevertheless, the strength in the response in differential gene expression in sporulating wild type cells varied between different experiments (Fig. 2I,J,K). Fig. 2L displays all data points of the comprehensive data set from Fig. 2A-K with a Simprof cluster-specific color coding.

**Fig. 2.**
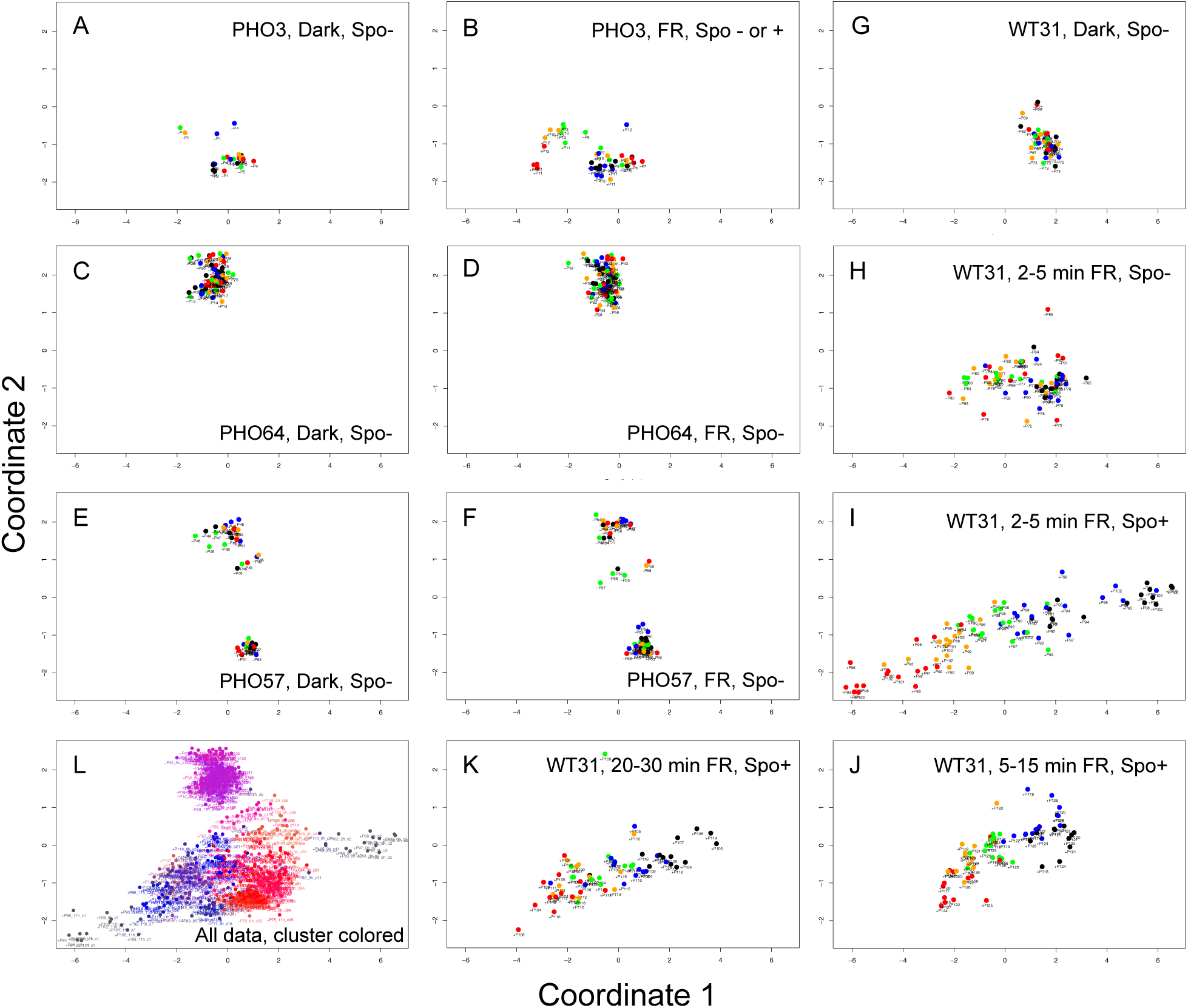
Multidimensional scaling analysis of expression patterns of mutants and wild type. Expression data of the experiments listed in Table 1 were pooled, normalized, analyzed by multidimensional scaling, and plotted to the same scale for each indicated class of cells separately. The color code of data points in panels A-K refers to the time at which the sample was taken (0h, black; 2h, blue; 6h, green; 8h, yellow; 11h, red). Panel L shows all data points of panels A-K, with colors according to their respective Simprof cluster ID number encoded by a rainbow spectrum. Abbreviations: FR, far-red stimulated; Spo-, Spo+, plasmodium had not sporulated or sporulated on the next day of the experiment, respectively. A character string of +P118_2h_c5, for example, states that the sample was taken from plasmodium number P118 at 2h after the start of the experiment, that the gene expression pattern of the sample was assigned to Simprof cluster number 5 and that the plasmodium had sporulated (+, sporulated; -, not sporulated) on the next day of the experiment (see Materials and Methods). These character strings (except those in Fig. 1) uniquely refer to the trajectories listed in Table 2 and SI Table 2 and also identify the respective sample in the heat maps of the comprehensive data set (SI Figs. 2,3).

Figure 3 shows typical single cell trajectories of mutant and wild type as picked from the data set of Fig. 2. Trajectories were all drawn to the same scale but arbitrarily arranged in the plane while keeping the orientation relative to the x and y axes of the MDS plot. According to the Simprof cluster ID number displayed adjacent to each data point there are significant changes in the gene expression patterns of successive time points within a trajectory even in unstimulated, not sporulating cells (Fig. 3). The trajectories of sporulating cells clearly moved from left to right in the PHO3 mutant (Fig. 3C) and in the wild type (Fig. 3J) while the response of stimulated but not sporulating cells of the two genotypes was less directed (Fig. 3B,I). Note that for the sporulating wild-type three very long, still typical trajectories have been selected for display while most of the trajectories of sporulating wild-type cells resembled the shorter and less straightly directed ones of Fig. 3 J. For the full set of single cell trajectories see SI File Package 1.

**Fig. 3.**
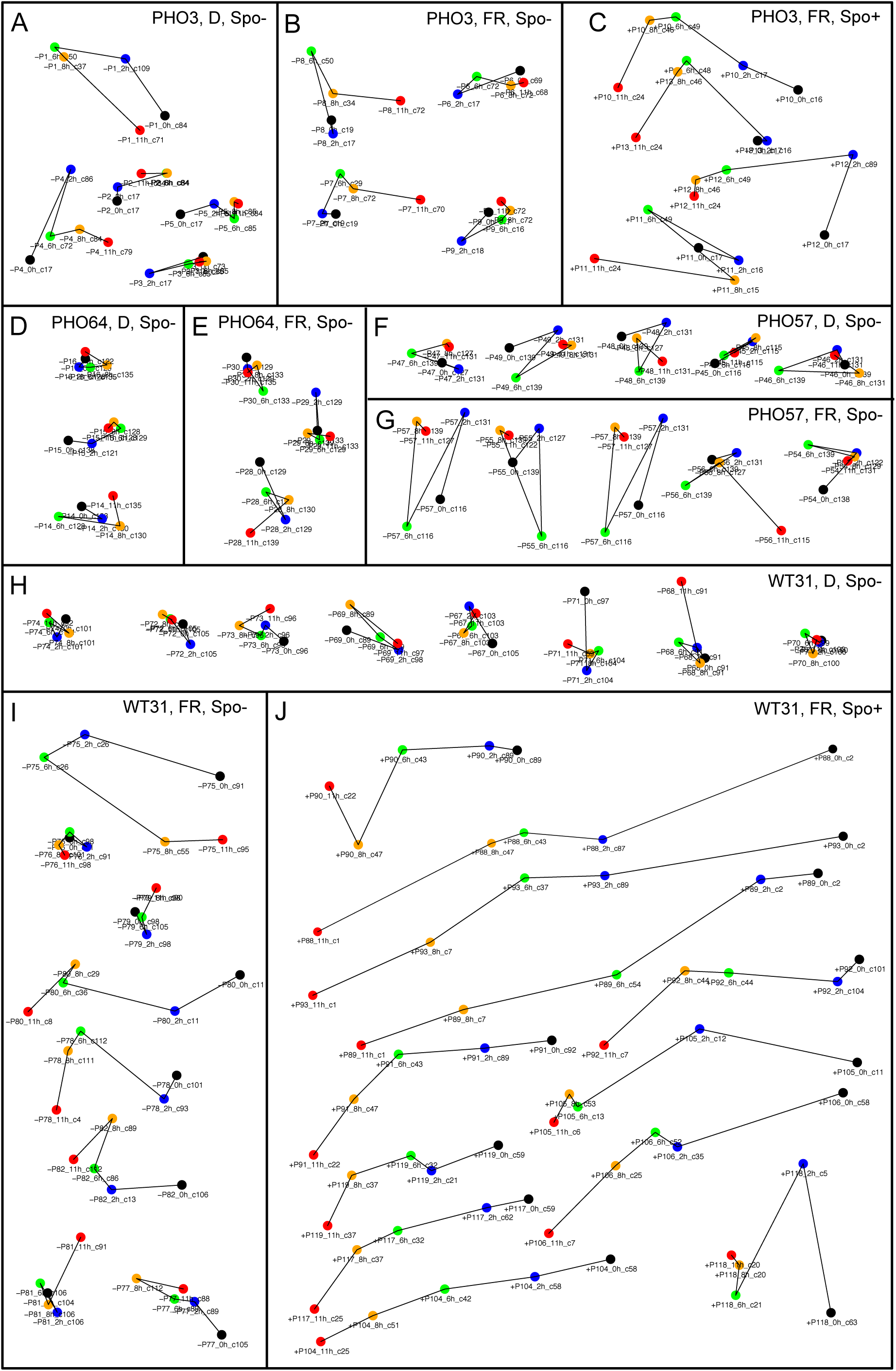
Typical single cell trajectories of mutant and wild-type cells picked from the data set displayed in Fig. 2. Tracks were arbitrarily arranged relative to each other in the plane of the panels while the relative orientation of the tracks with respect to the x-and y-axes of the multidimensional scaling plots was maintained. Data points were labelled as described in the legend of Fig. 2. Abbreviations: FR, far-red stimulated; Spo-, Spo+, plasmodium had not sporulated or sporulated on the next day of the experiment, respectively.

### 3.1 Constructing Petri nets from single cell trajectories of gene expression

As in most of the MDS trajectories of wild type and mutants, cells change from one Simprof cluster to another between successive time points, we have defined trajectories through the subsequent states of gene expression of each cell according to the Simprof cluster ID number (SI Table 2). These states of gene expression and the changes between them can be easily represented in the form of a Petri net (Werthmann and Marwan 2017). This is schematically shown for a simplified example considering the expression strength of two genes at three time points (Fig. 4). Each significant cluster is represented as a place (drawn as circle) of a Petri net and the change between two clusters is represented by a transition (drawn as rectangle, connecting two places via directed edges (arcs)). The current gene expression state of a cell is indicated by a token that resides in one of the places. The Petri net is marked with a single token at any time, as a cell can only be in one state of gene expression at the same time. If a cell transits from one state to another, a transition moves the token to a downstream place as indicated by the directed edges. Keeping this terminology, we discriminate between *transit* (the change between gene expression states of a living cell) and *transition* (the Petri net node). Because of the structure of the Petri net, tokens can neither be produced nor destroyed (for details see legend of Fig. 4). Depending on the number of genes considered by clustering, each place may represent a complex pattern of gene expression in the form of a vector of positive real numbers. Accordingly, the number of places and the overall structure of the Petri net constructed from a data set depend on how the gene expression patterns change with time but also on the clustering procedure that evaluates these changes.

**Fig. 4.**
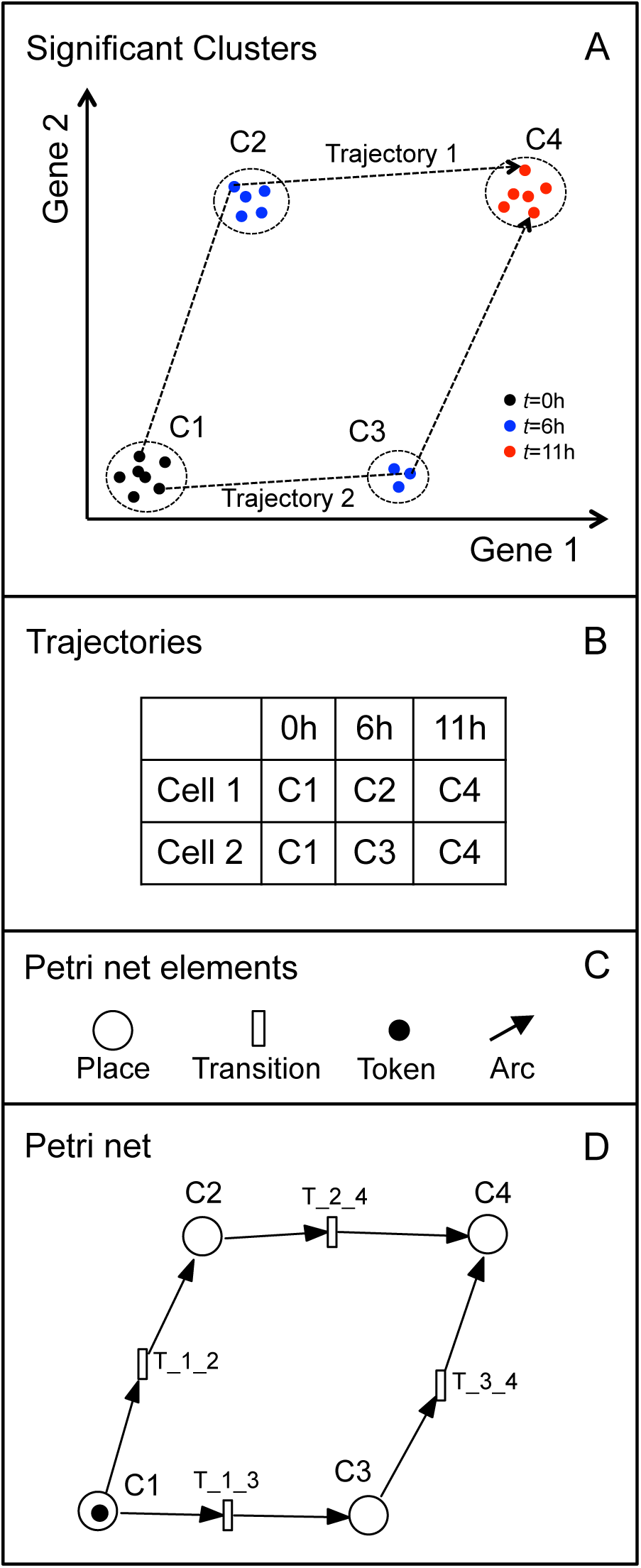
Automatic construction of Petri nets from single cell trajectories. (A) The state of gene expression of each cell at any given time point was classified by its assignment to a significant cluster as determined with the help of the Simprof algorithm. For the sake of simplicity, the scheme considers two genes and three time points only. (B) The trajectory of a single cell was defined by the subsequent states of gene expression through which the cell proceeded. (C) Elements of a Petri net in their graphic representation. (D) Each state of gene expression as characterized by its Simprof cluster assignment was represented by a place and each transit between two states was represented by a transition linking the two places with arcs, indicating the direction in time in which the transit occurred. The current state of gene expression of a cell is indicated by a token marking the respective place. The Petri net is marked with one single token, as the cell can be only in one state of gene expression at the same time. Petri nets accordingly constructed from any set of single cell trajectories consist of one single P-invariant: because each transition has exactly one ingoing and one outgoing arc and because all arc weights are one, tokens can neither be produced nor destroyed and the state of gene expression remains unequivocally defined.

Extended stochastic Petri nets (Heiner et al. 2009) were constructed automatically from single cell trajectories of gene expression and were encoded implicitly considering the transits between successive states (Simprof significant clusters). These implicit representations were translated into stochastic Petri nets encoded as ANDL files (Abstract Net Description Language (Heiner et al. 2013)) that were imported into Snoopy (Heiner et al. 2012; Rohr et al. 2010) and graphically displayed by running the Sugiyama (Sugiyama et al. 1981) or the planarization layout algorithm implemented in Snoopy (Heiner et al. 2012). For details see SI Computational Methods. Assigning class numbers to user-defined subsets of the data set (Table 1) allowed the construction of a set of Petri nets each representing the entire data set, only one subset of the data, or an arbitrary combination of subsets.

### 3.2 Structural features of the Petri nets

To demonstrate some basic structural features of the obtained Petri nets and to discuss the information they reveal, let us consider a small net of sporulating cells. Figure 5 shows Petri nets from two different experiments in which all cells sporulated.

**Fig. 5.**
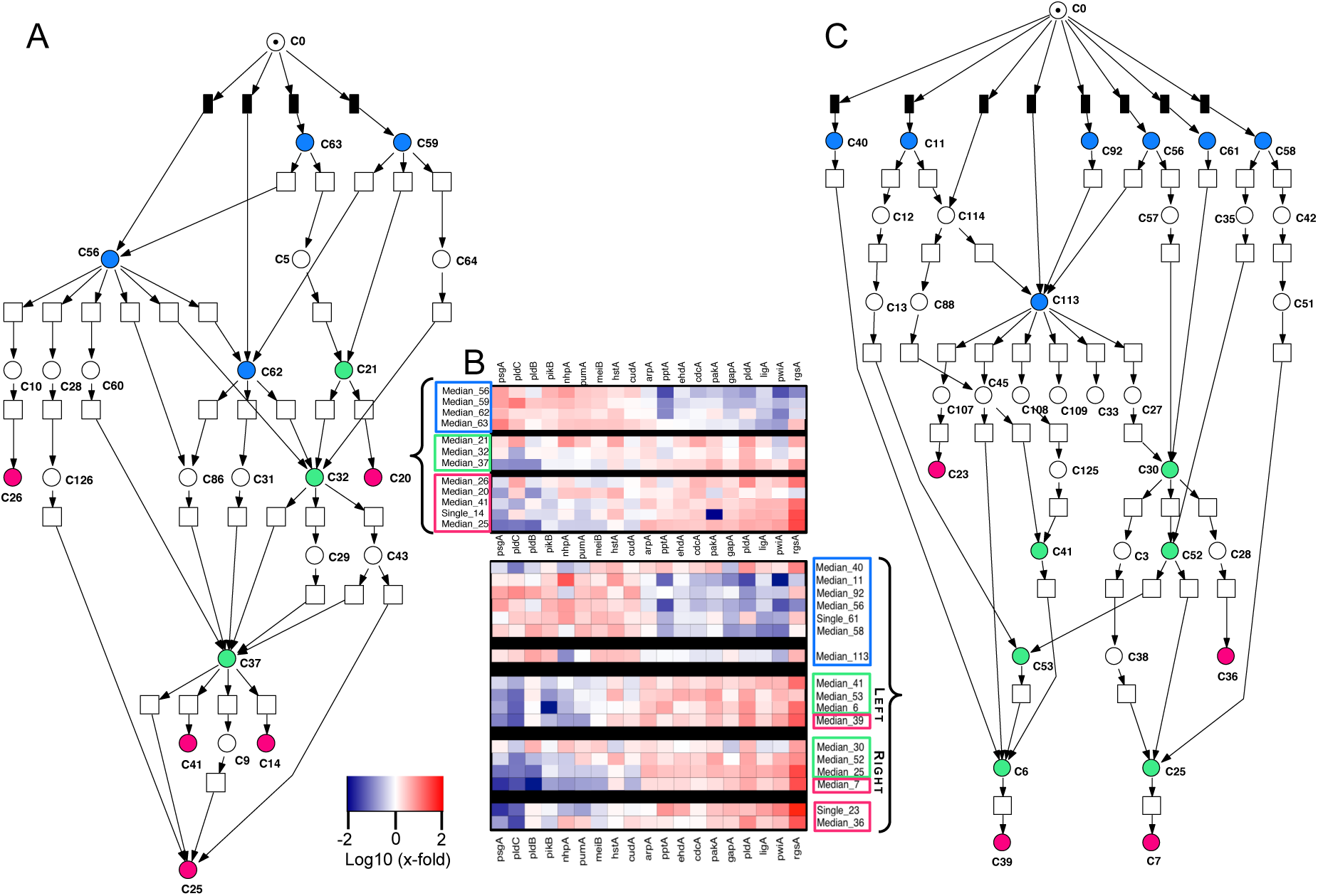
Petri nets of wild type plasmodial cells sporulating in response to a saturating far-red light pulse. Petri nets were constructed from data sets of cells that were starved for six (A) or seven (C) days, corresponding to subsets 11 and 10 in Table 1, respectively. Color coding of places: starting point of a trajectory, blue; moderately or highly connected places, green; sink places, carmine. Immediate transitions are filled in black. (B) Heat map showing the gene expression patterns of the Simprof clusters (median or single values), with cluster ID numbers indicated. The colors of the rectangles refer to the coloring of the places.

#### Starting points of trajectories

The starting point of a trajectory is referred to as the state of gene expression in which the cell resided at the start of the experiment (0h) and is defined by the corresponding Simprof cluster ID number. To graphically highlight places referring to starting points of trajectories and to automatically create the initial marking of Petri nets in stochastic simulations, immediate transitions (Heiner et al. 2009) (filled in black) all connected to the same pre-place (C0) were used. Place C0 is introduced for technical reasons only. It does not refer to any cluster of gene expression and it is marked with one token for initialization purposes. At the start of a simulation, each immediate transition has the chance to deliver the token from place C0 into its downstream place (*i.e.* into the post-place of the immediate transition) in which at least one cell trajectory started. The relative firing propensity of each immediate transition was automatically set proportional to the number of trajectories that started in the state represented by the place the immediate transition is connected to. This generates an according distribution of initially marked Petri nets when stochastic simulations are repeatedly run, *e.g.* for finding sink places (see below) by simulative model checking.

#### Source places

Places from which tokens can only flow out are called source places. When the immediate transitions used for initialization are deleted (or optionally not even implemented during automatic Petri net construction, see SI Computational Methods), at least some of the places that represent trajectory starting points are source places. Note that trajectories do not necessarily start in a source place (e.g. C56 in Fig. 5A). For a formal treatment of source places, sink places, T-invariants, *etc.* see (Heiner 2009).

#### Sink places

Places that have only ingoing arcs are called sink places as tokens cannot flow out again. When the only token of a net is trapped in a sink place, the net has reached a terminal state (*dead lock* in Petri net terminology) where none of the transitions can fire any more. Sink places accordingly mark the end of trajectories. However, trajectories did not necessarily end in a sink place (Table 2). Even if a trajectory ends in a sink place, the true end of the cell’s trajectory may not have been reached yet. Extending the time series may hence well turn the apparent terminal state into a transient state. Hence the sink places are referred to as structural entities of the Petri nets presented here but do not necessarily correspond to terminal physiological states of a cell (in case such terminal states should even exist). However, we observed that there are sink places where a number of arcs converge suggesting that these places represent attractor states or at least metastable intermediates (see Discussion).

#### T-invariants

A T-invariant defines a reaction cycle that brings the sub-system, which is covered by this T-invariant back to the state from which it started. There is not a single T-invariant in the nets of Fig. 5 indicating that the flow of tokens is entirely directed through irreversible steps. T-invariants however were observed in the Petri nets of wild type cells that did not sporulate (Fig. 6A,B) and in the Petri nets of non-sporulating mutants.

**Fig. 6.**
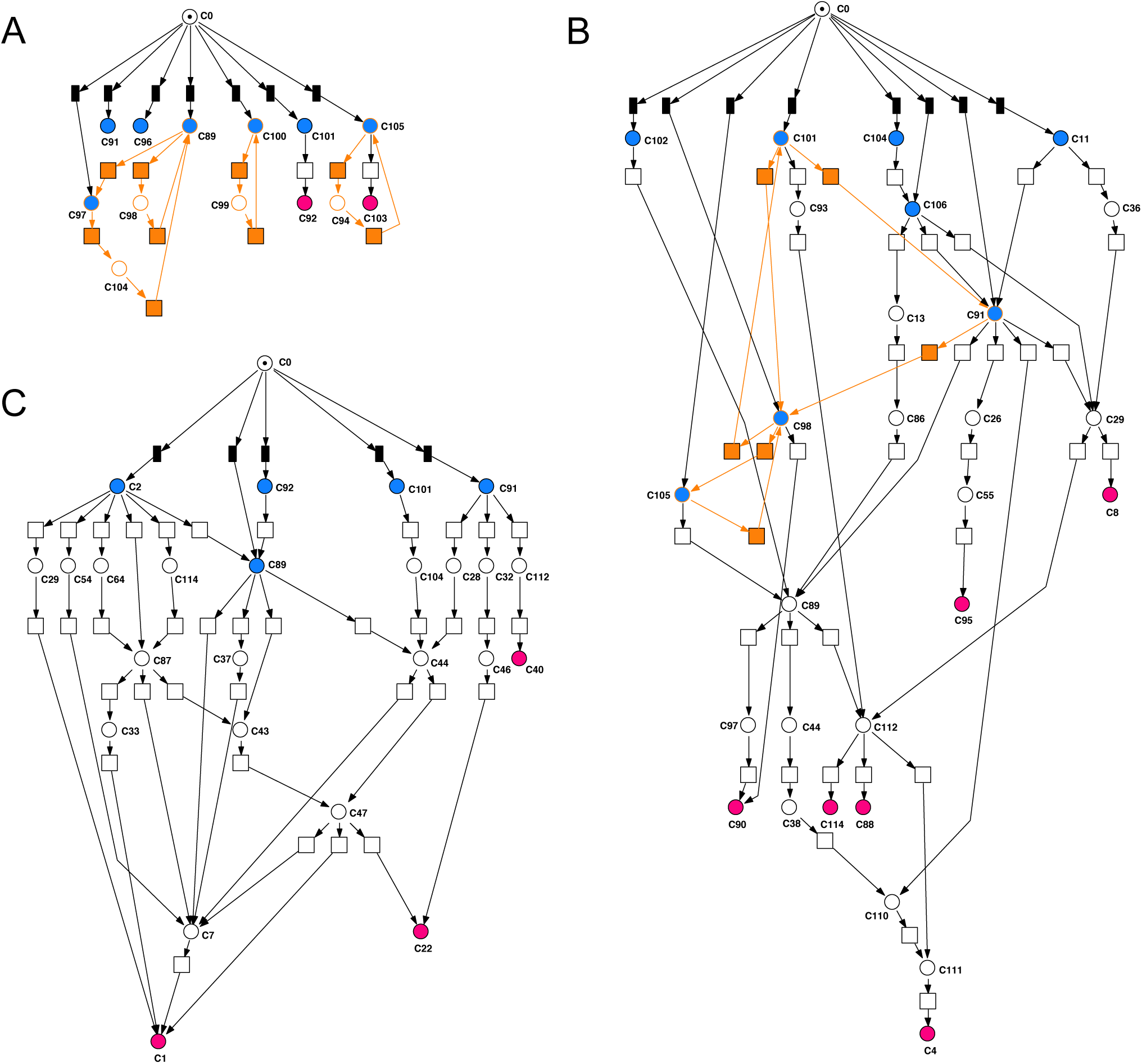
Response of wild type cells to threshold stimulation with far-red light. Petri nets of unstimulated cells (A), of stimulated, yet not sporulating cells (B), and of cells that sporulated in response to the stimulus (C) were constructed separately from the transits that occurred in classes 7, 8, and 9 in Table 1, respectively. Color coding: starting point of a trajectory, blue; sink places, carmine; T-invariants, orange. Immediate transitions are filled in black.

#### Highly connected places

We found places that were connected to multiple transitions by in-and/or out-going arcs (e.g. C56, C62, C37, or C25 in Fig. 5A). These highly connected places obviously represent states that are relatively likely. Accordingly, they may represent metastable intermediates or even states of an attractor. In the Petri nets displayed in this work, those sink places that are not highly connected might nevertheless correspond to attractor states which are infrequently reached.

The Sugiyama algorithm (Sugiyama et al. 1981), which was used for automatic layout highlights the directedness of processes and facilitates the visual identification of sink places. (Note that source places and sink places should be identified computationally through (simulative) model checking which certainly is independent on any graphical representation.) The Sugiyama layout also nicely arranges the highly connected places relative to each other, which is especially helpful considering the small nets like those of Figs. 5 and 6.

The two Petri nets of Fig. 5 revealed important characteristics of far-red light-induced sporulation.

1. Trajectories start in different places (some of which are connected to each other) that represent significantly different states of gene expression.
2. Following light induction, sporulation occurs through parallel, branched and partly interconnected pathways that end in sink places (marked in carmine red in Fig. 5A,C) representing significantly different states of gene expression.
3. Sporulation did occur through intermediate states (e.g. C21, C32 in Fig. 5A and C113 in Fig. 5C) where genes are already up-regulated while down-regulation had not yet occurred.
4. There are highly connected places which may represent meta-stable or stable states of the dynamic system and correspondingly depressions or (local) minima (valleys) of the Waddington landscape (see Discussion).

Even when considering the low number of 19 differentially regulated genes, pathways to sporulation were variable. Single cell trajectories (Table 2, SI Table 2) in addition indicate that there is no rigid timing for switching from one cluster to the next. The cell that switched directly from its initial state C40 to the committed state C6 at 2h after the far-red pulse and then to sink place C39 at 6h in which it stayed until the 11h time point (Fig. 5C, SI Table 2) seems to be special as compared to the others. Possibly, this cell would have sporulated spontaneously even without light stimulus, which occasionally happens.

### 3.3 The response to threshold stimulation: comparing sporulating and not sporulating cells

While the Petri nets of Fig. 5 reveal alternative routes to sporulation through transient states of gene expression, the tipping points at which the irreversible commitment to sporulation occurs are not obvious. To identify commitment points by Petri net construction, we have analyzed the plasmodial response to stimulation at threshold, *i*.*e*. to a far-red light pulse which is so short that only approximately 50% of the plasmodial population sporulated. Sporulation of an individual plasmodial cell in response to a threshold stimulus is all or none (see Introduction). To have a dark control included, we have evaluated three groups of wild type cells: Cells that were not stimulated (class 7), cells that were stimulated but did not sporulate (class 8) and cells that sporulated in response to a stimulus (class 9). Although the Petri nets constructed for each of the three groups in part use the same places, we displayed the three nets separately in order to reveal the structural differences between them (later on the nets are merged; see below).

The net of the dark controls (Fig. 6A) consists of 13 places (except C0), considerably less than the nets of light-stimulated cells (Table 1). It contains one source place, two sink places and two places which are not connected to any stochastic transition, resulting from the trajectories of cells that stayed in the same gene expression state throughout the experiment. In addition, the net contains places that are part of four T-invariants, indicating cyclic reaction paths. Places and T-invariants may appear as isolated entities simply because the number of cells analyzed was not sufficiently high and possible transits were missed out accordingly. Nevertheless, it suggests that unstimulated wild type cells do have some spontaneous activity in changing between states of gene expression (Fig. 6).

Cells that have been stimulated but did not sporulate (Fig. 6B, Table 2) reveal a Petri net with considerably more places (28, except C0) connected by 39 stochastic transitions. The net has 8 places in which trajectories started, 4 of them are source places (after removal of the immediate transitions). The net contains 3 T-invariants (shown in orange), indicating possible reaction cycles that lead back to previous states. There are six sink places that differ from the sink places of sporulating cells but also from the places representing starting points of trajectories. The observation that a number of pre-places of sink places are not part of any T-invariant suggests that the cells that have been stimulated but did not take the decision to sporulate and hence remained in the plasmodial state instead, nevertheless did not return to the original, unstimulated states, at least not within the observation period of 11h.

The Petri net constructed from the trajectories of plasmodial cells sporulating in response to threshold stimulation (Fig. 6C) consists of 24 places (except C0) and 36 transitions, four source places and three sink places. It contains not a single T-invariant, indicating that the cells moved straightforward to sporulation through irreversible steps proceeding through alternative pathways. This is consistent with the straightforward, yet variable tracks seen in the multi-dimensional scaling plots (Fig. 3J). Nine of 16 trajectories of sporulating cells started in C2 (Table 2) taking different directions as indicated by the six arcs that connect C2 to different downstream places. This obvious difference between this net and the net of stimulated but non-sporulatmg cells (Fig. 6B), in which C2 does not even occur, suggests that cells which were in C2 had a higher probability to sporulate in response to the weak (near threshold) far-red light stimulus than cells with another starting point of their trajectory. Interestingly, the expression levels of the up-regulated genes including the mRNA of the transcription factor homolog *arpA* is very low in C2 as compared to the other initial states (Fig. 7) suggesting that *arpA* expression might be associated with enhanced propensity to enter sporulation. The structural features of the Petri nets of wild-type cells of classes 10 and 11 sporulating in response to saturating stimulation (Fig. 5 A,C) are indeed similar to those described for the net of Class 9 (Fig. 6C): The cells started at different places, there are no T-Invariants and the token proceeds irreversibly towards different highly connected places or to sink places downstream of these places. An evaluation of the three nets of sporulating plasmodia with respect to common places will be performed by combining the three nets into one coherent net (see below).

**Fig. 7.**
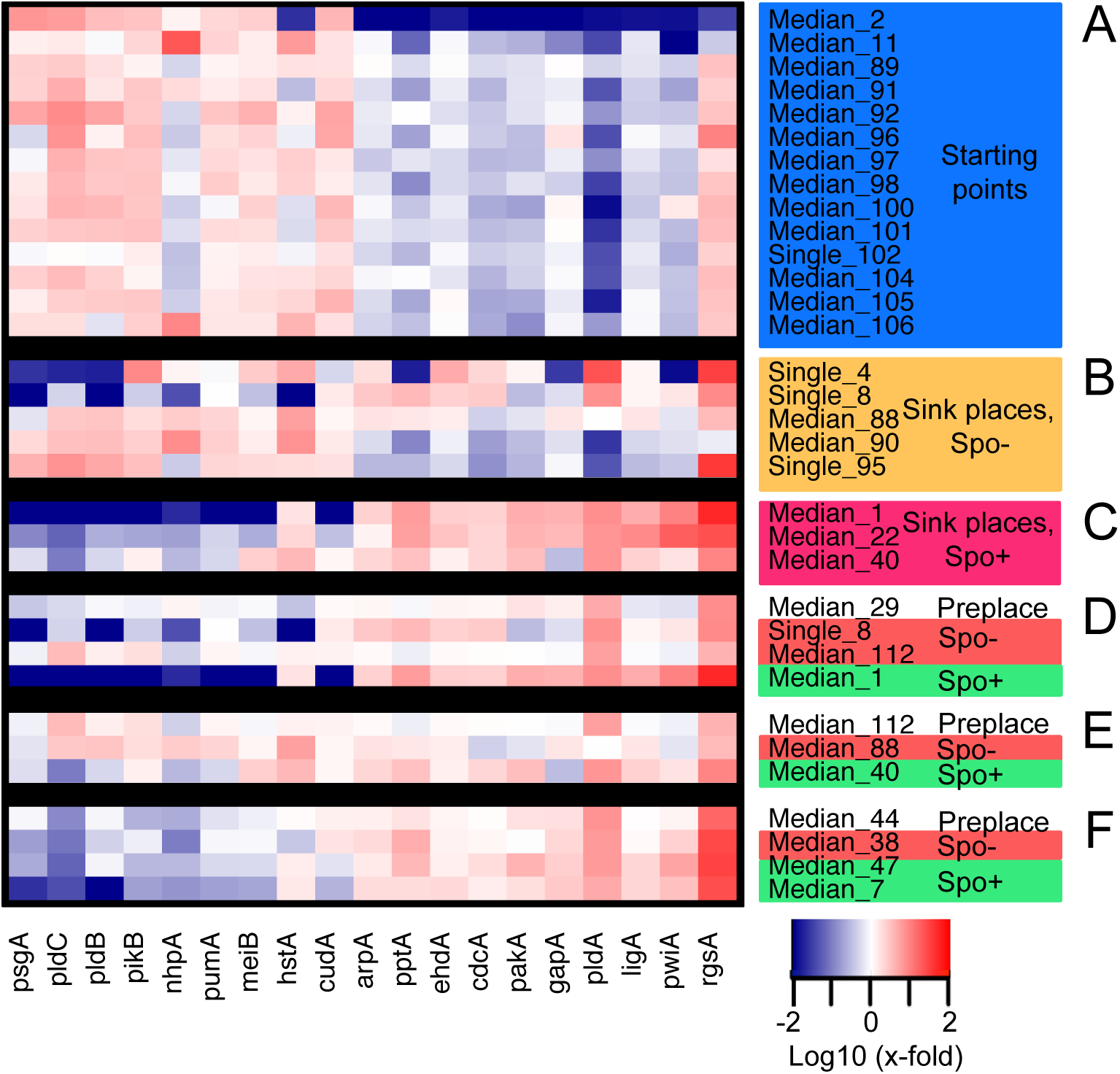
Gene expression patterns of wild type cells in response to threshold stimulation with far-red light. Expression patterns are displayed for a selected set of places of Petri nets in Figs. 6 and 8 referring to starting points of a trajectory (A) or to sink places found in not sporulating (B) or in sporulating (C) cells. (D-F) Expression patterns referring to places that mark branching points linked to transitions irreversibly leading to a sporulated (Spo+) or to a not sporulated state (Spo-) as identified in Fig. 8.

### 3.4 Transitions associated with the commitment to sporulation

To identify transitions that are associated with the irreversible commitment to sporulation, the nets of subsets 7,8,9 were merged into one Petri net (Fig. 8). The merging algorithm of the Pascal program generated a name for each transition that contained the number of each subset (class) of cells in which the corresponding transit occurred. This allowed to colorize the transitions according to their occurrence in the respective data class using the *Search node* option in Snoopy. As expected, T-invariants mainly occurred through transitions specific for not sporulating cells (FR light stimulated and dark controls, classes 7 and 8, see above), indicating the possibility to cycle between states. Not-sporulating and sporulating cells shared a number of places that were not trajectory starting points (e.g. C44, C89, C112, C114 *etc.* in Fig. 8). In the planarization layout of the net however, transitions corresponding to not sporulating (orange transitions) or to sporulating cells (green transitions) where predominantly located in different, island-like regions of the net (Fig. 8B), highlighting those places that were specific for sporulating or for not sporulating cells.

**Fig. 8.**
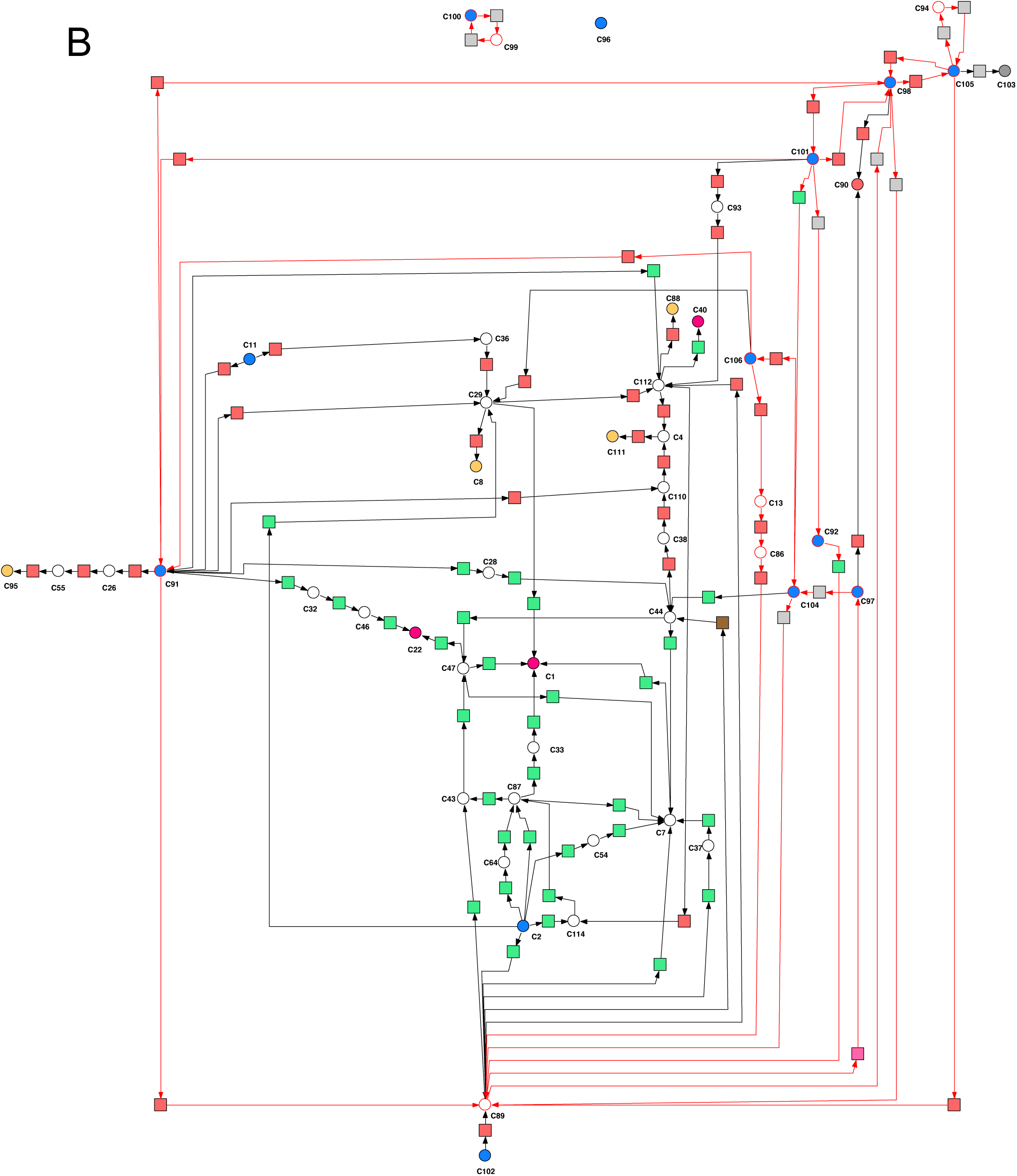
Petri net-based identification of expression patterns associated with the commitment to sporulation. The three Petri nets of Fig. 6 were combined into one coherent net and transitions were colored according to whether transits occurred in unstimulated cells, in cells that did not sporulate, or in cells that sporulated in response to stimulation. The Sugiyama layout (A) facilitates the visual identification of branching points with respect to the developmental decision. Branching points are defined as places which are directly linked to at least one transition that irreversibly leads to sporulation and simultaneously to at least one alternative transition that irreversibly leads to a not sporulated state. Expression patterns corresponding to the three branching points (C29, C112, and C44) in the net are shown in Fig. 7. (B) Layout of the same net with the Planarization algorithm yields an alternative graphical representation in which places directly linked by a transition are drawn close to each other while arc crossing is largely avoided. T-invariants in (A) and (B) are indicated by arcs drawn in orange.

The Sugiyama layout facilitates the visual identification of tipping points in terms of transitions that irreversibly led to the sporulated or to the not sporulated state. These transitions are certainly not part of any T-invariant. As seen in Fig. 8A, the transition from C29 to C1 led to sporulation while the transition from C29 to C8 led to the not sporulated state of the plasmodium. Other places directly connected to downstream transitions representing competing tipping points were C112 and C44. The transition from C89 to C37 caused the exit from a T-invariant to enter the irreversible path to the sporulated state (C1). The gene expression patterns corresponding to places directly upstream or downstream of tipping point transitions are shown in Fig. 7 (D-F).

**Fig. 8A.**
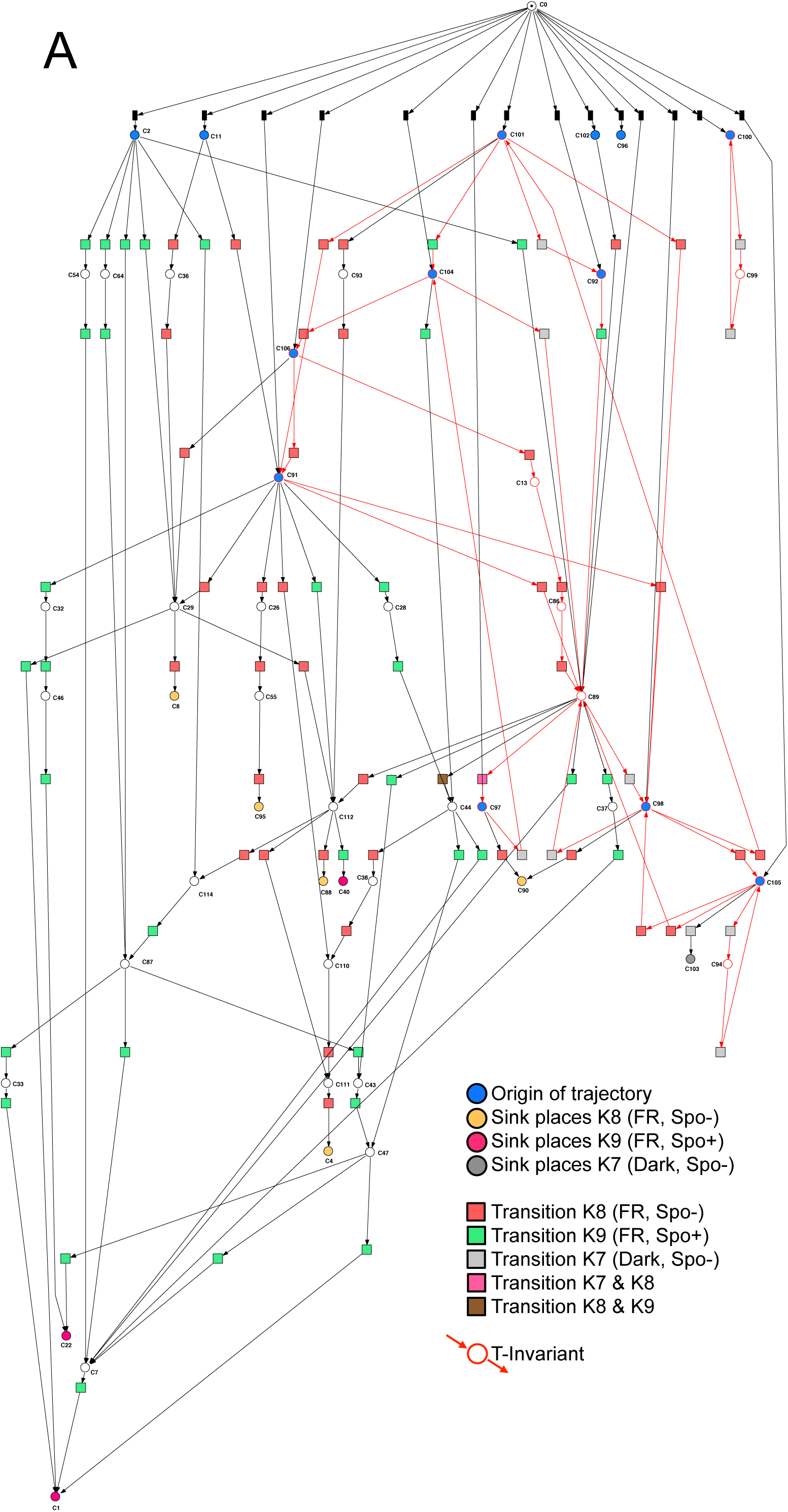
See next page for legend.

As we have shown previously, differential gene regulation does occur in response to far-red light well before commitment (Hoffmann et al. 2012; Rätzel and Marwan 2015). The expression data in Fig. 7 together with the Petri net in Fig. 8 suggest that an elevated level of the up-regulated genes is a pre-requisite for commitment while commitment itself seems to be associated with the extensive down-regulation of genes. The expression patterns suggest that *rgsA* and *pldA* are strongly expressed even before commitment (Fig. 7 D-F). But otherwise we did not recognize any obvious order of differential regulation.

### 3.5 Petri nets of mutants with reduced propensity to sporulate

We have analyzed three mutant strains with reduced (PHO3) or strongly reduced (PHO64, PHO57) propensity to sporulate. When PHO3 cells were exposed to a pulse of far-red light which is highly saturating in causing sporulation in the wild type, approximately one out of two PHO3 plasmodia sporulated in an all-or-none response. In contrast, PHO64 and PHO57 cells were locked in the plasmodial state and did not sporulate. None of the two strains showed obvious differences in the gene expression patterns caused by the light stimulus. However, there were differences in the expression level of some genes in PHO64 and even more pronounced in PHO57 (Fig. 1) with trajectories indicating that cells visited different clusters during the time window of the experiment (SI Table 2). As there was no obvious response in gene expression to far-red light stimulation, data of stimulated and unstimulated cells were pooled for Petri net construction and the transitions were colored to discriminate transits specific for either group of cells. The Petri net of PH064 cells (Fig. 9) contains 21 places connected by 59 stochastic transitions confirming that cells spontaneously switched between different states of gene expression. The net is covered with 1809 T-invariants indicating extensive possibilities to cycle between states. There were hardly any places that might be specific for stimulated or unstimulated cells and many transitions occurred in both classes (red transitions in Fig. 9). The heat map displayed for a selection of places indicates that there are genes that clearly change between high and low states. Some genes were abundantly expressed like in sporulating wild type cells while *rgsA* was low throughout.

PHO57, similar to PHO64 cells, neither sporulated nor showed this mutant any obvious changes in the gene expression patterns in response to far-red light stimulation. In contrast to PHO64 however, the PHO57 data gave two separate Petri nets (Fig. 10), obviously corresponding to the two well-separated clouds of data points in the MDS plots of Fig. 2E,F.

**Fig. 9.**
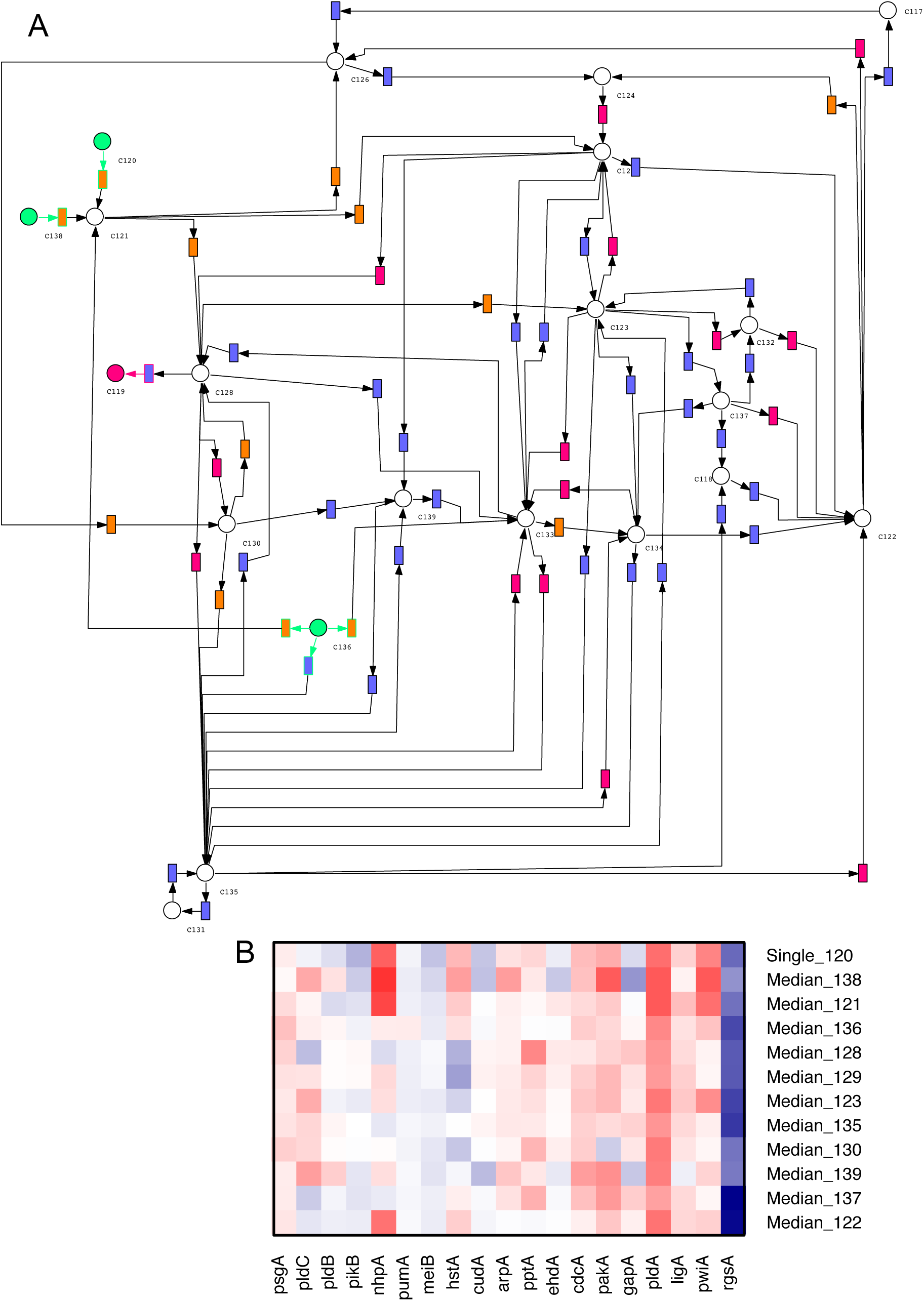
Petri net constructed from trajectories of the non-sporulating mutant PHO64. (A) The net is composed of transitions that were specific for transits in unstimulated cells (orange, class 3), for transits in stimulated cells (blue, class 4) and for transits that occurred both, in unstimulated and in stimulated cells (carmine). The net is covered with 1809 T-invariants. Source places are shown in green, the only sink place in carmine. (B) Expression patterns corresponding to selected places of the net displayed in approximate order of their location from the left to the right side of the net.

In total, the two disconnected nets of PHO57 are composed of 24 places connected by 45 stochastic transitions and covered with 31 T-invariants. Each of the two disconnected nets is composed of transitions from stimulated cells and dark controls, suggesting that the cells, whether stimulated or not, resided in two distinct macro-states while the probability of switching between the two states was at least so low that it did not occur in the limited number of samples analyzed. The heat map (Fig. 10) displayed for a set of selected places reveals substantial differences between the two nets in the expression of a set of genes *(pakA, pldA, rgsA, etc.)* while the expression levels of *nhpA* and *pwiA* drastically varied in both of the two nets.

**Fig. 10.**
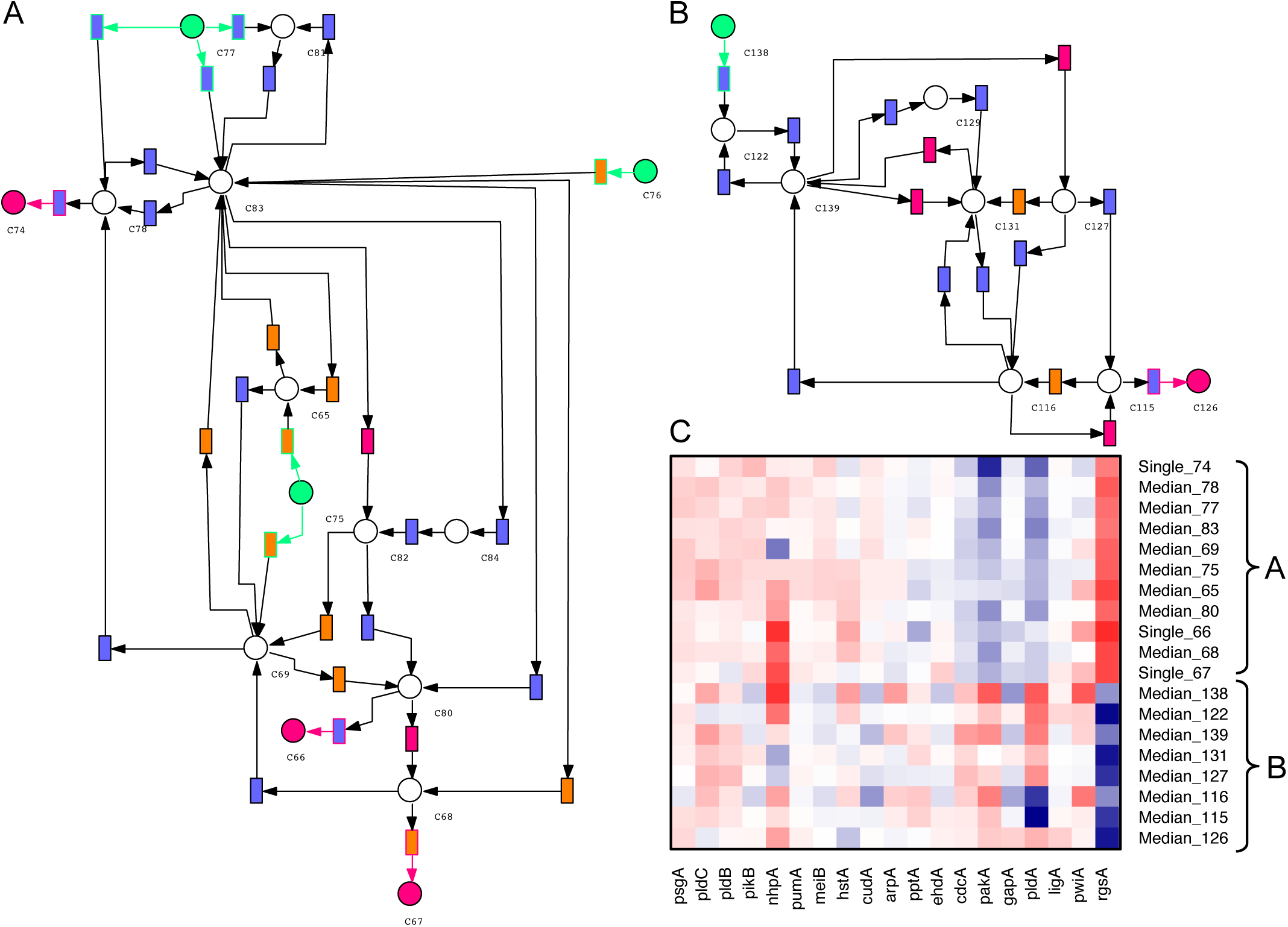
Petri nets constructed from trajectories of the non-sporulating mutant PHO57. (A, B) Trajectories of unstimulated and stimulated cells revealed two disconnected Petri nets. Transitions are colored according to the occurrence of transits in unstimulated cells (K5, orange), in stimulated cells (K6, blue) or in both (K5 and K6, carmine). The two nets are covered with a total of 31 T-invariants. Source places are marked in green and sink places in carmine. (C) Gene expression patterns corresponding to selected places of the nets in (A) and (B).

The Petri net of PHO3 cells (Fig. 11) which is composed of far-red stimulated sporulating and not sporulating cells along with not sporulating dark controls (26 places, 38 stochastic transitions) differed from the nets of PHO57 and PHO64 cells in being covered by only 3 T-invariants that were specific for not sporulating cells while the response to sporulating cells was directed. Except for *rgsA* which was relatively high throughout, the expression patterns were qualitatively similar to those of sporulating or not sporulating wild type cells, respectively while some genes even in the dark controls seemed to be spontaneously up-or down-regulated (heat map in Fig. 11).

**Fig. 11.**
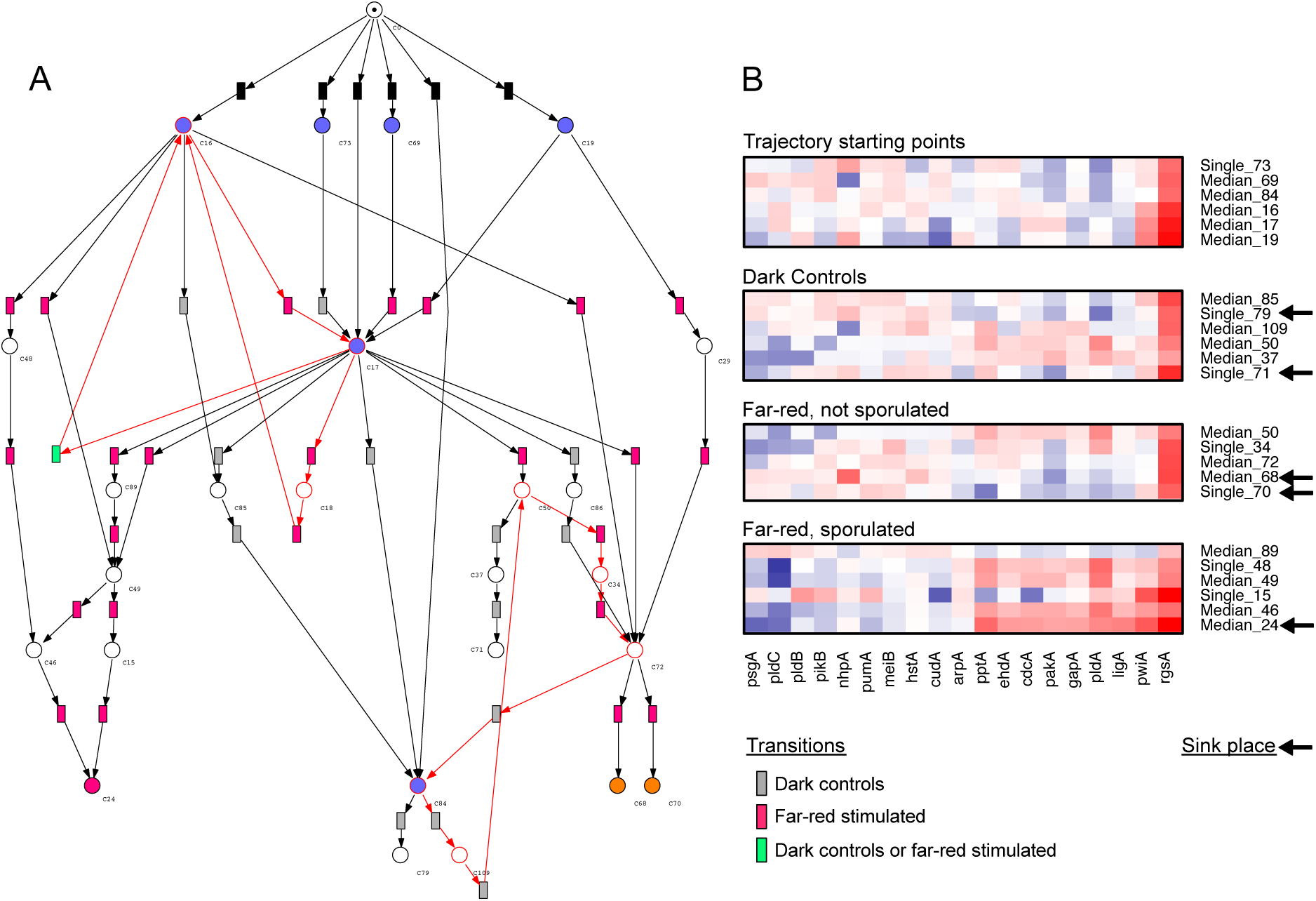
Petri net constructed from trajectories of PH03 cells. (A) Transitions are colored according to the occurrence of transits in unstimulated cells (class 1, grey), in stimulated cells (class 2, carmine) or in both (class 1 and class 2, green). Places referring to the starting point of a trajectory are marked in blue, the sink place of sporulated cells in carmine and the sink places of stimulated, yet not sporulated cells in orange. Immediate transitions are filled in black. The net contains three T-invariants (marked in red) that were specific for not sporulated cells. (B) Gene expression patterns corresponding to selected places referring to starting points of trajectories, to unstimulated cells, or to cells that did not or did sporulate in response to stimulation.

### 3.6 Petri net composed of data from wild type and mutants

To structurally compare the nets obtained for wild type and mutants, all data subsets of Table 1 were combined into one comprehensive Petri net. The transitions were colored according to the class number assigned to the corresponding subset of the data in which the transit occurred. Automatic layout revealed one coherent Petri net (Fig. 12) and one isolated place (C96) which resulted from one single, unstimulated wild type cell that stayed in the same gene expression state during the time of the experiment.

The planarization layout algorithm distributed the net into two rectangular areas with one place (C126) connecting the two rectangles through wild type-and PH064-specific transitions.

**Fig. 12.**
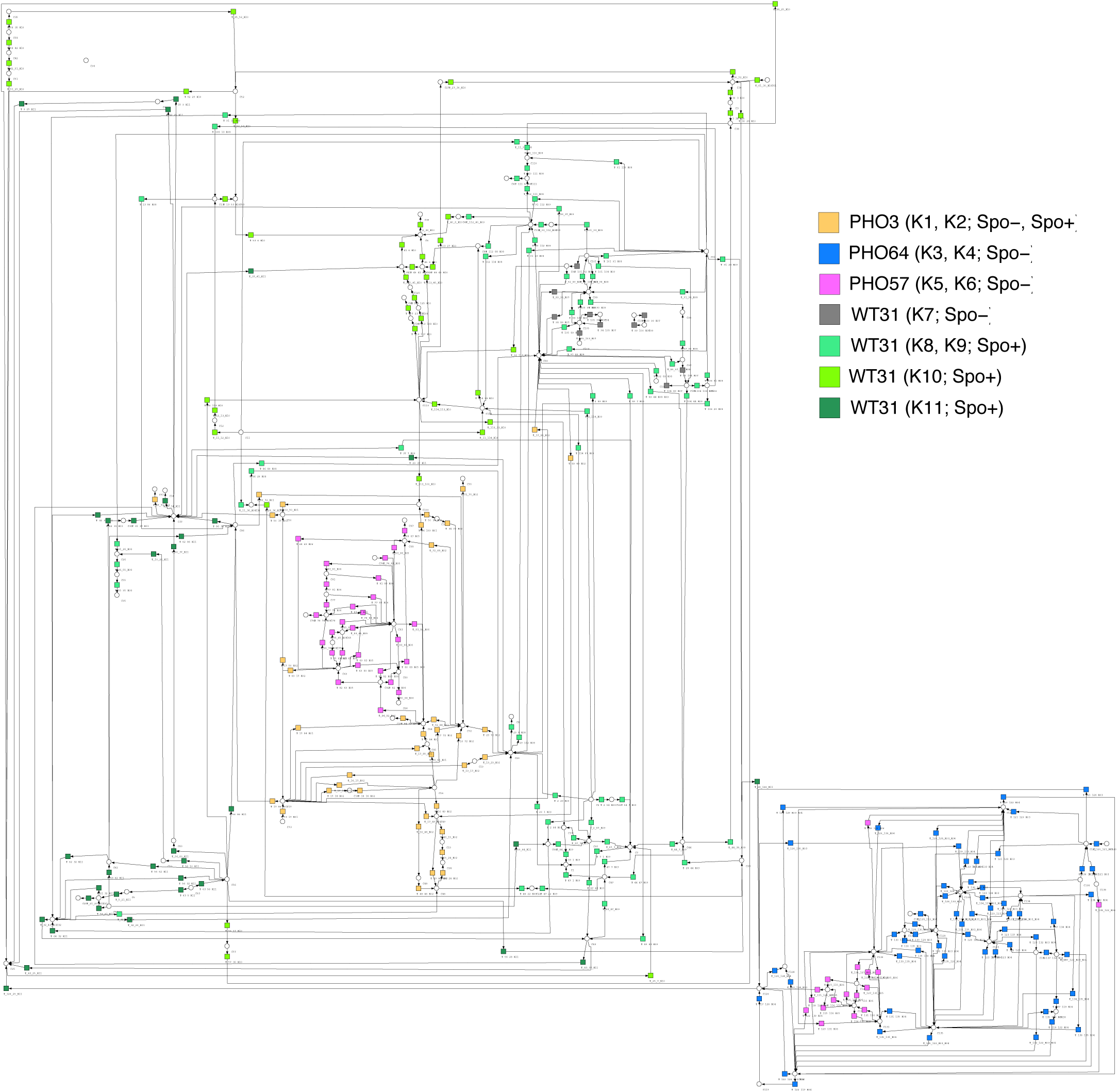
Petri nets constructed from the complete comprehensive data set as displayed in Table 1. Transitions were colored according to the data subset in which the corresponding transits occurred as indicated by the inset. In preparing the figure, the subsequent coloring of the transitions was recorded in the form of a flip-book movie (SI File Package 1) to reveal potential overlapping assignment of transitions to more than one subset.

The subnet of the small rectangle is composed of places and transitions specific for PHO64 and PHO57 with transitions in blue and pink, respectively that form distinct, strain-specific islands connected by few common places. The remaining set of PHO57-specific places and transitions is located in the center of the subnet of the large rectangle, connected to the rest of the net via few PHO3-specific transitions. For wild type and mutants, transitions of equal color are clearly not randomly distributed over the net, indicating that transitions of each color preferentially connect certain subgroups of places. Accordingly, the net reveals which gene expression states (places) and which transitions are preferentially specific for wild type and mutants as well as for sporulating and not sporulating cells, respectively. The net also identifies places that are shared between wild type and mutants and suggests transitions that may be specific for a certain subset (Table 1).

## 4 Discussion

### 4.1 Variability in the cellular response of light-induced sporulation

Before discussing the structural features of the obtained Petri nets, we will consider sources of variability of molecular events involved in light-induced sporulation. Over a period of many years we have noticed that sporulation-competent plasmodial wild type cells of any given strain that were grown and starved under standardized experimental conditions (Starostzik and Marwan 1998) to a certain degree do vary in light sensitivity for the induction of sporulation. This is presumably due to small changes in the growth dynamics of the culture. There is also a certain degree of variability in the morphology of fruiting bodies between individual plasmodia of one culture batch and between the plasmodia of different culture batches. On top of this variability, light sensitivity and the sporulation propensity increases with the period of starvation (5d, 6d, 7d *etc.*). Because of this variability we have deliberately constructed Petri nets from data sets from different experiments obtained under different experimental conditions to see whether the developmental program of sporulation is variable to a certain extent at the level of differential gene expression. As expected, we found Petri nets with different structural details and with a number of places that occurred in only one of the three nets constructed for the subsets of sporulating cells. When the three data sets were combined however, automatic construction revealed one coherent Petri net, indicating that there was also a number of shared places between the three nets, so there is variation and overlap.

### 4.2 Petri nets reveal dynamic routes to commitment and differentiation

We have analyzed gene expression time series of wild type and mutant cells in response to a stimulus pulse of far-red light. The vigorous cytoplasmic streaming which continually mixes the large volume of the cytoplasm is expected to dampen stochasticity of signaling and of the gene expression events which would result in molecular noise within the small cytoplasmic volume of a typical mononucleate mammalian cell. Accordingly, changes in the gene expression pattern measured in the *P. polycephalum* plasmodium and the resulting single cell trajectories are expected to reflect dynamic phenomena within un-stimulated and stimulated cells rather than stochastic fluctuations in the number of mRNA molecules transcribed from any given individual gene.

We identified significantly different states in gene expression with the help of the *Simprof* cluster algorithm (Clarke et al. 2008), assigned a Petri net place to each cluster of states, and determined the transits between any of the successive states for each single cell trajectory. Each transit between two states as discretized by clustering was considered as firing event of a corresponding transition connecting the two places representing the two clusters of states. In this way, single cell trajectories were determined from the data set of expression patterns and then directly translated into a corresponding Petri net. The wiring diagram of the Petri net displays the ways on which significantly different states may be interconverted into each other. The approach obeys to Occam’s razor or to the principle of parsimony in assuming that the states as defined by being a member of a significant cluster are identical unless it has been proven that they are different. Discretization of states by clustering was performed by choosing a statistical significance level of α=0.05 (5% error probability) for the Simprof algorithm ensuring that obviously different gene expression patterns were assigned to clusters with different ID numbers. Accordingly, Petri net places represent gene expression states that differ highly significantly from each other.

The purpose of this work was to explore and demonstrate the potential power of the Petri net approach to visualize, analyze, model, simulate, and understand the dynamics of states of cellular regulation during commitment and differentiation. At this time, we do not aim at establishing structural details of the constructed Petri nets nor do we draw any final conclusion regarding system dynamics or mechanisms which the nets might suggest. Instead, we report and discuss the properties of the nets in their present form, bearing in mind that they were constructed from a data set of still limited size and with a limited resolution in time. In future experiments we need to analyze more cells, more genes, and more time points to work out and establish structural details of the nets. Accordingly, currently observed differences in trajectories between groups of cells (e.g. light-exposed versus dark controls) that translate into structural features of a Petri net might emerge or disappear. With more data analyzed, one might observe that places that represent seemingly parallel ways in connecting two places through different intermediate states turn out to be sequential steps within the same pathway (Fig. 13A). This also applies to seemingly alternative pathways where one directly connects two places while the other involves an additional place as intermediate state (Fig. 13B). This caveat appears to be most relevant for places that have one ingoing and one outgoing arc rather than for highly connected places.

**Fig. 13.**
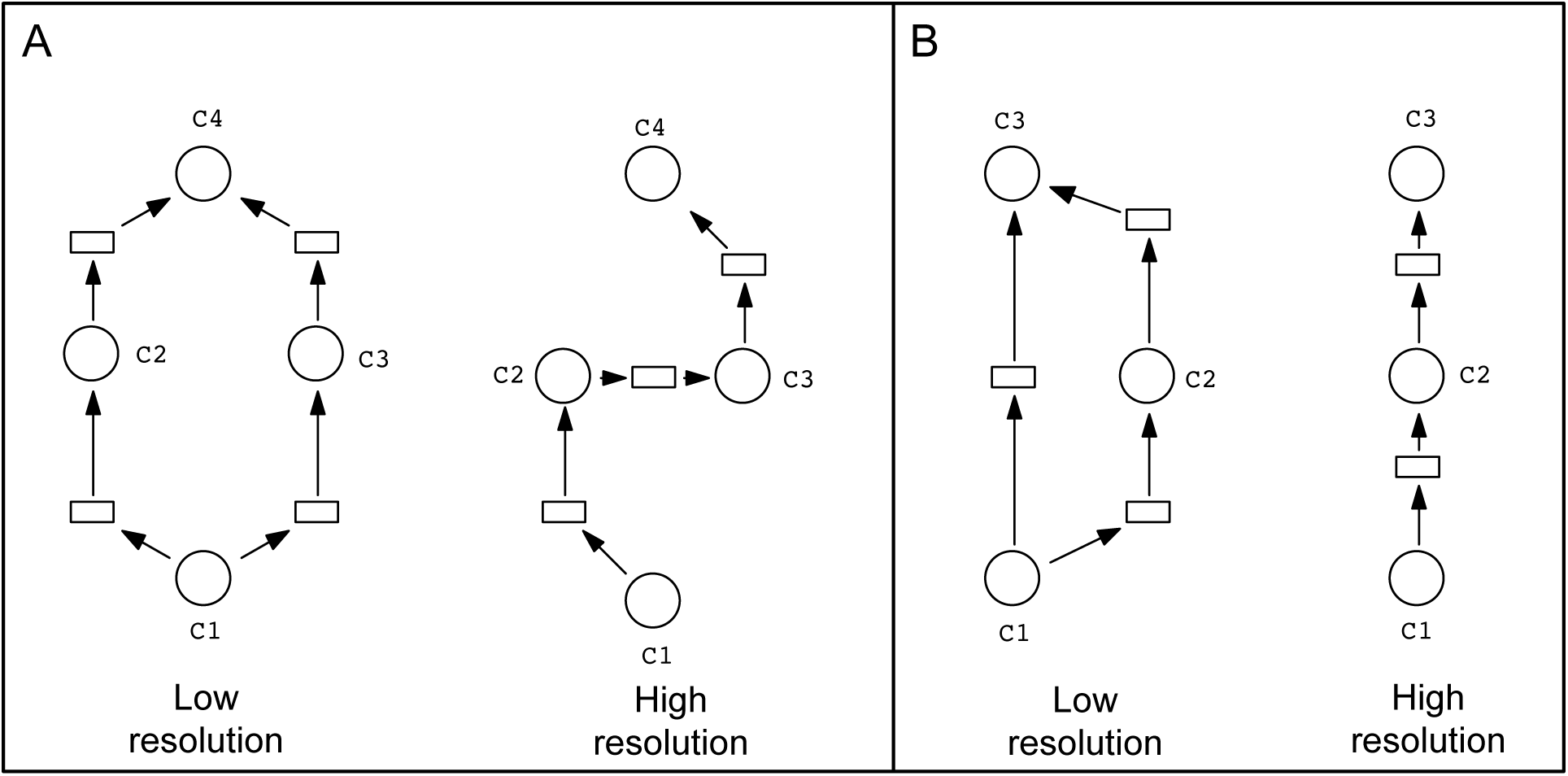
Network motifs indicating the causal sequence of events may become better resolved to approach the correct structure with growing sample size and higher time resolution. (A) and (B) display examples as discussed.

Despite these words of caution, the results obtained in this study do suggest that there are indeed numerous qualitatively different pathways leading from the proliferative, plasmodial state to sporulation and that these pathways can be identified through the Petri net approach. Different pathways are even directly evident from those trajectories that lead to sporulation starting from highly different states when being stimulated with light.

There are two extreme scenarios of how the decision for light-induced sporulation might be taken: In one scenario, activation of the phytochrome receptor triggers a rigid, well-defined sequence of molecular events that leads to commitment and sporulation. In the alternative scenario of a dynamic system, activation of phytochrome triggers molecular events that destabilize the previously stable proliferative plasmodial state so that the cell sporulates by entering a new stable state. In this case, the molecular events that occur upon the transit from one stable state to the other are variable and less predictable. This variability could for example be just the result of the causal system dynamics where the quantitative interplay of a certain set of molecular components may drastically change in response to small changes in the side conditions and accordingly different trajectories emerge that nevertheless lead to the same attractor, the stable state of the sporulated cell. Another or additional source of variation could for example be the different internal state of the cell (e.g. the phase of the cell cycle) in the moment of phytochrome activation which then leads to an adaptive self-reorganization of molecular events. The highly variable developmental trajectories of *P. polycephalum* plasmodia to commitment and sporulation (Rätzel and Marwan 2015) do not support the scenario of a rigid sequence of events.

### 4.3 Probing the Waddington landscape with Petri nets

We suppose that cell fate decisions and developmental reprogramming of cells are controled by the quasi-potential landscape of a dynamic system which we simply call *Waddington landscape.* In addition to the cumulative evidence mentioned in the Introduction, this assumption is justified by the discovery that a system of interconnected switches forms multiple stable states through so-called attractors (Bornholdt and Kauffman 2019; Kauffman 1969). The assumption is evenly justified by the thorough work on bifurcations and other dynamic phenomena that emerge from well-understood kinetic mechanisms of cellular regulation (Ferrell Jr 2012; Tyson et al. 2003; Tyson et al. 2019). In principle, one may obtain a graphical representation of a Waddington landscape by projecting the *n*-dimensional state space of a system of *n* components (or genes) into a two-dimensional plane. Plotting the negative logarithm of the probability that the system is in a certain state as the third dimension then gives a landscape in which mountain tops correspond to unstable and hence unlikely states and valleys correspond to likely, stable states or to so-called attractors ((Huang et al. 2009; Zhou and Huang 2011) and references therein). It has been shown by several groups that such quasi-potential landscapes may result from the dynamic behavior of well-known molecular kinetic mechanisms ((Ferrell Jr 2012; Wu et al. 2017) and references therein).

Deriving the quasi-potential landscape of cellular reprogramming by computation is challenging because it would require knowledge of mechanistic details, kinetic rate constants, and concentrations which all can affect the dynamic behavior of a network (Endres 2012; Rowland et al. 2012; Varusai et al. 2015; Vilar et al. 2003; Zhou et al. 2012). This knowledge is sketchy even for mammalian cells (Zhou et al. 2012) and it remains unknown how the wiring diagram of a gene regulatory network, the gene expression noise, and signal transduction altogether shape the attractor landscape and determine the developmental trajectories of a cell (Wu et al. 2017). Accordingly, we directly observed the cellular behavior and explored a data-driven, equation-independent modeling approach based on Petri nets.

Some structural features of the small networks of sporulating cells as they appear in the Sugiyama layout (Figs. 5,6), suggest that the Waddington landscape indeed can be probed by Petri net construction. Two significantly different gene expression patterns of a cell represented by two places of a Petri net do represent two correspondingly distinct points in the plane of the Waddington landscape (see above). The token, sequentially marking Petri net places may be accordingly viewed as the marble in Waddington’s paradigm rolling through the landscape being trapped by basins of attraction. The probability (propensity) that a certain transition fires to move the token from one place to the next would then presumably be related to the quasi-potential that drives the corresponding change in gene expression.

We have shown that a cell can start at different places when receiving an inductive light stimulus. It then proceeds in the direction of significantly different terminal states represented by sink places that are reached with different probability as indicated by the experimentally observed single cell trajectories. We propose that the highly connected places represent states that are likely to occur and therefore do occur frequently. This suggests that these states indicate meta-stable intermediates or even terminal states that reside in a basin of attraction.

T-invariants indicate cyclic transits between states and these states might belong to a common basin of attraction. The token can escape with a certain probability depending on the firing propensity of connecting transitions that do not belong to the T-invariant.

Although we have constructed stochastic Petri nets, the firing propensities of the stochastic transitions have not been adjusted to fit the experimental data and therefore do not refer to the true probability for switching between states. Running stochastic simulation nevertheless allowed the identification of sink places by simulative model checking which is computationally less expensive for large nets than structural analysis.

The Petri net structure turned out to be helpful in revealing critical transitions that start a sequence of firing events that is essentially irreversible. This allows the identification of tipping points, the molecular alterations associated with them, and their genetic control.

Moreover, the Petri net approach allows the characterization of complex phenotypic alterations caused by mutations. Through the islands of equally colored transitions that specifically refer to cells of different genotype, the Petri net of Fig. 12 clearly shows how sporulation propensity-affecting mutations shift the probability of gene expression states as compared to the wild type. Shifts in gene expression are also evident by the creation of new T-invariants that identify potential mutation-induced attractors.

The sporulation-suppressing mutations in PH064 and PH057 lock the cells in the proliferative state and suppress differentiation. In PH03, the proliferative plasmodial state is clearly stabilized at the expense of the propensity to differentiate. The formation of new T-invariants and of genotype-specific islands of transitions as observed in these mutants is especially interesting as it supports the idea that mutations or other oncogenic alterations act through the generation of cancer attractors (Brock and Huang 2017; Huang et al. 2009).

### 4.4 Reconstruction of differential gene regulation

The places of the Petri nets as we have used them implicitly represent states of gene expression, here expressed as the median of the expression values of the considered genes within each Simprof cluster. These values can be discretized and together with the Petri net structure one can obtain feasible sequences of state-and reaction vectors as input to reconstruct the underlying gene regulatory network. Including genetic information, network reconstruction (Durzinsky et al. 2013; Durzinsky et al. 2011; Marwan et al. 2008) can then reveal causal interactions that are represented again in the form of a Petri net. This however is beyond the scope of this article.

### 4.5 Conclusions and outlook

We have described a Petri net-based approach to disentangle the complex dynamics of differential gene expression in the context of cell commitment and differentiation. We further demonstrated how Petri nets can be used to reveal and analyze complex phenotypes caused by mutation and proposed that the combination with appropriate network inference techniques should ultimately allow to reconstruct the underlying gene regulatory network.

As the formal approach described here does neither depend on equations nor on prior knowledge of the internal structure of the analyzed system, it seems likely that it is accordingly applicable to the analysis of other complex systems or processes if time series data on sets of observables can be obtained. In analogy to the effect of gene mutations which we demonstrated, one might be able to see how factors of interest can change attractor landscapes or identify tipping points that depend on such factors. This may perhaps be of interest for the study of certain phenomena in ecology, economy, or in the social sciences.

Of particular interest might be the application of the Petri net approach to the system level analysis of pathogenesis, disease progression, or the response to drug treatment. In cases where underlying mechanisms are complex and poorly understood, disentangling the complexity of individual responses and the identification of potential tipping points based on a set of clinical parameters followed as a function of time may perhaps be helpful even if the causal connections of the measured parameters to the phenomenon of interest are not even established. The Petri net approach might nevertheless be able to translate a collection of clinical data into a visual and executable model to predict the response of individuals based on their personal history.

## 6 Author’s Contributions

VR, BW and JS performed single cell time-series experiments and gene expression analyses to generate the data sets. BW and VR evaluated results, explored suitable statistical analysis methods, and laid the ground for the data evaluation pipeline. MH supervised part of the experimental work and contributed a data management system. WM conceived of and supervised the study, performed computational analyses, developed the data analysis pipeline including the automated generation of Petri nets, evaluated results, and wrote the paper. All authors read and approved the final version of the manuscript.

## 8 Acknowledgments

We thank Mrs. Bärbel Lorenz and Mrs. Bianca Mertens for excellent technical assistance.

## 9 Supplementary Information

SI File Package 1: Supplemental Tables, Figures

Supplementary Computational Methods

## Supplementary Computational Methods

### Automated generation of Petri nets from single cell trajectories

Petri nets were automatically constructed from the transits as determined by the R script (see the Materials and Methods section of the main manuscript for details) stored in a Petri net description vector (PNDV) in an implicit format and exported from R as a text file. To do so, the transits that occurred between each pair of states were cumulated to give one corresponding transition in the Petri net. In the implicit representation by the PNDV (SI Fig. 4A), each transition was encoded by three integers, the number of the Simprof cluster characterizing the state of the cell *before* the transit, the number of the Simprof cluster characterizing the state of the cell *after* the transit, and the number of transits (events) that occurred in the data set.

In the R script of the data analysis pipeline, the user could define subsets of data within the data set to allow comparison of e.g. stimulated and unstimulated cells or of wild type and mutants *etc.* The algorithm then exported one Petri net encompassing the entire data set and a set of separate Petri nets, one for each subset.

### Generation of ANDL Files

ANDL files (SI Fig. 5) were generated with a program written in Turbo Pascal which was run in the DOSBox under Mac OS X (https://www.dosbox.com). The program reads the implicit representations of the Petri nets exported as text files from R (SI Fig. 4) and generated two ANDL files from each text file to encode one stochastic Petri net without and one with initialization through immediate transitions as explained in the main text of this article.

Accordingly, the Pascal program generates a pair of ANDL files encoding Petri nets encompassing the entire data set and a pair of ANDL files for each user-defined subset of the data set as exported from R. To evaluate how changes in conditions impact the structure of a Petri net, the Pascal program allows to generate ANDL files of Petri nets by combining Petri net files of subsets of data as specified by the user. The program uses the Petri net description vector exported from R and generates transition names for the ANDL file that indicate in which subset(s) of the data the particular transits occurred and, optionally, how often the transit happened. After import into Snoopy, transitions can then be colorized in a subset-specific manner using the *Search nodes* function of Snoopy (see below). The source code and the compiled form of the Pascal program that can be directly run in the DOSBox are provided as part of the Supplementary File Package 4 (Petri net construction kit).

**SI Fig. 4.**
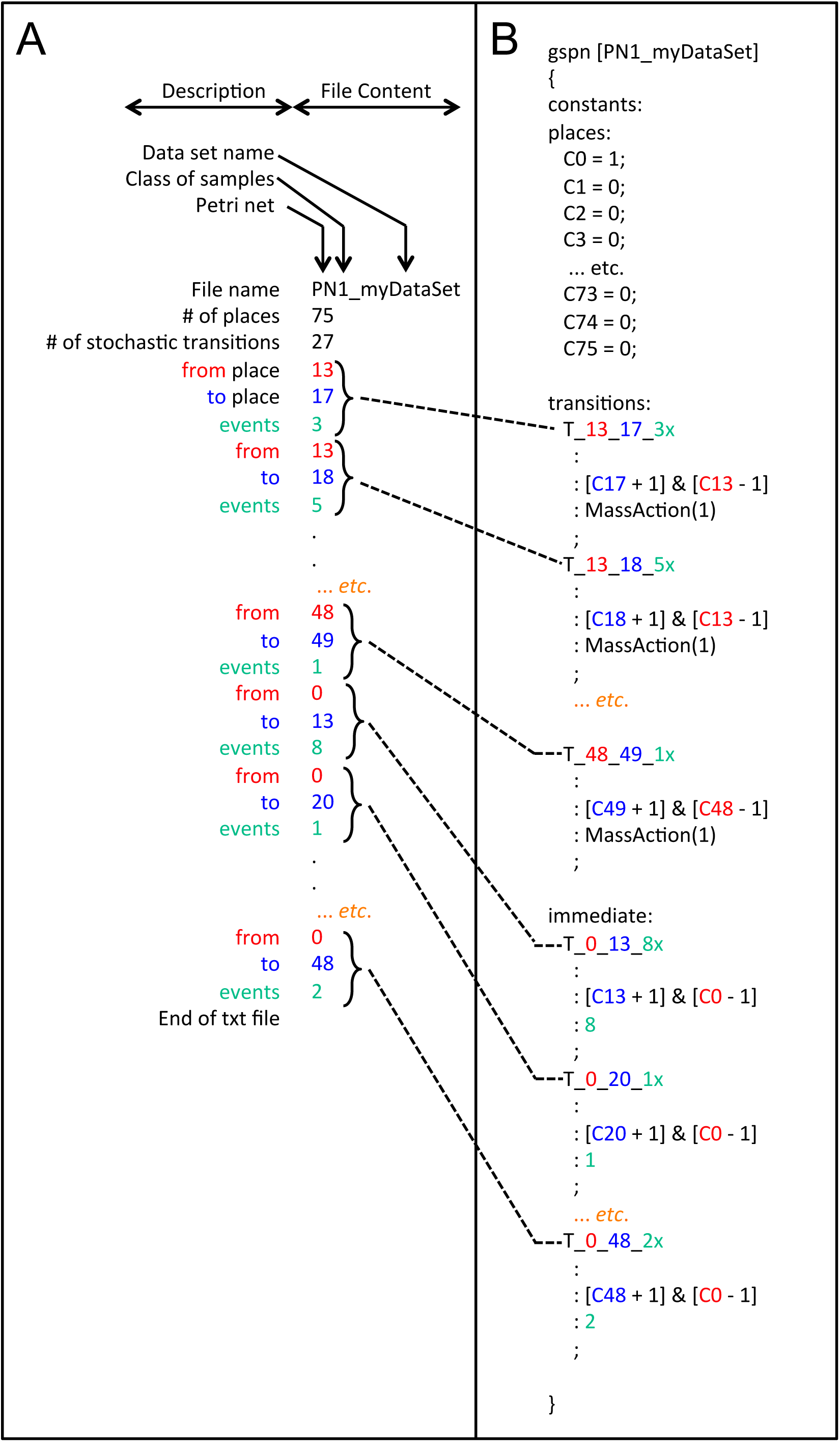
Structure of the Petri net description vector (PNDV) as exported in the form of a text file from R and its conversion into an ANDL file. The ANDL file is generated by a Pascal program and subsequently imported into Snoopy for graphical display of the Petri net. (A) Example of the content of a PNDV-containing file encoding the Petri net and description of its elements. Note that the description is not part of the text file. The first line contains the name of the file (without extension) as automatically generated in R based on the name of the analyzed data set, the second line contains the total number of places (except C0) and the third line contains the total number of *stochastic* transitions of the Petri net. The rest of the file is composed of a sequence of lines in defined order, each line containing one integer number to encode information about stochastic or immediate transitions. Stochastic transitions are listed before immediate transitions. Each transition, stochastic or immediate, is encoded by three subsequent integers specifying (1) the pre-place and (2) the post-place of the transition and (3) the number of transits between Simprof clusters that occurred in the set or subset of the data. These three integers are then used by the Pascal program to generate the name of the transition, its connectivity, and the firing propensity in the case of immediate transitions as part of an ANDL file as shown in (B). In this study, the firing propensity of stochastic transitions was automatically set to 1 for generation of the ANDL file.

**SI Fig. 5.**
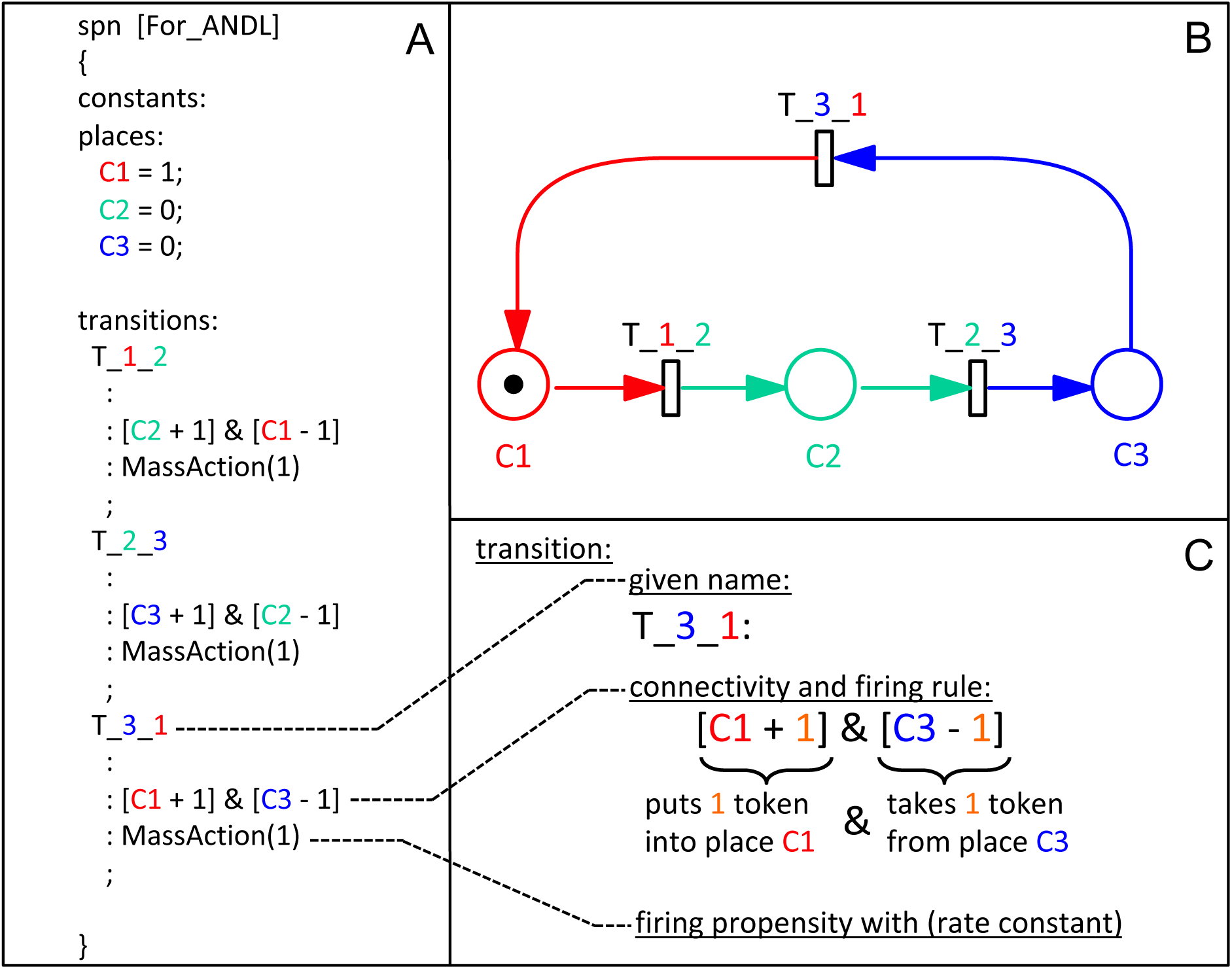
ANDL syntax used for encoding stochastic Petri nets for the subsequent import into Snoopy. (A) Complete code of the ANDL file encoding the stochastic Petri net graphically displayed in (B). (C) Explanation of the syntax for encoding each stochastic transition of the Petri net with (1) the name of the transition as defined by the user, (2) the connectivity of the transition to places of the net through in-and out-going arcs including the arc weights, *i.e*. the number of tokens taken from the pre-place(s) and delivered into the post-place(s) upon firing of the transition, and (3) the stochastic firing propensity of the transition in the form of the rate constant referring to mass action kinetics. Note that Petri nets that are manually drawn in Snoopy can be exported as ANDL files which helps to get familiar with the syntax.

### Generation and graphical layout of stochastic Petri nets in Snoopy

Stochastic Petri nets were generated by importing a chosen ANDL file into Snoopy, As an ANDL file does not contain any information regarding the graphical presentation of the Petri net, graphical layout was done automatically with the help of the *Layout* function using the *Sugiyama* or the *Planarization* algorithm implemented in Snoopy. In order to highlight and to color transitions with respect to the occurrence of the corresponding transits in a given subset of the data, transitions were selected through string elements of their names using the *Search nodes* function. The selected transitions were then colorized using the *Edit selected elements* function that allows to adjust the graphical representation of all selected elements.

